# Remodeling Ca^2+^ dynamics by targeting a promising E-box containing G-quadruplex at *ORAI1* promoter in triple-negative breast cancer

**DOI:** 10.1101/2024.03.28.587164

**Authors:** Oishika Chatterjee, Jagannath Jana, Suman Panda, Anindya Dutta, Akshay Sharma, Suman Saurav, Rajender K. Motiani, Klaus Weisz, Subhrangsu Chatterjee

## Abstract

ORAI1 is an intrinsic component of store-operated calcium entry (SOCE) that strictly regulates Ca^2+^ influx in most non-excitable cells. ORAI1 has been extensively studied to have been overexpressed in various cancer phenotypes, and its signal transduction has been associated with oncotherapy resistance. There is extensive proteomic interaction of ORAI1 with other channels and effectors, resulting in various altered phenotypes. However, the transcription regulation of this gene is not well understood. We have found a putative G-quadruplex (G4) motif, *ORAI1-Pu*, in the upstream promoter region of the gene, having regulatory functions. High-resolution 3-D NMR structure elucidation suggests that *ORAI1-Pu* is a stable parallel-stranded G4, having an unusual 8-nt loop imparting dynamics without affecting the structural stability. The protruded loop further houses an E-box motif that provides a docking site for transcription factors like Zeb1. The G4 structure was also endogenously observed using Chromatin Immunoprecipitation (ChIP) with anti-G4 antibody (BG4) in the MDA-MB-231 cell line overexpressing *ORAI1*. Ligand-mediated stabilization suggested that the stabilized G4 represses transcription in cancer cell line MDA-MB-231. Downregulation of transcription further cascaded down to a decrease in Ca^2+^ entry by the SOCE pathway, as observed by Fura-2 confocal Ca^2+^ imaging.

## Introduction

Metastasis continues to remain the leading cause of death for cancer patients, accounting for 90% of cases of the disease, despite notable advancements in diagnostic and therapeutic techniques. Among the most crucial regulators of proliferation and metastasis is the ubiquitous second messenger Calcium ions (Ca^2+^).^1^ Ca^2+^ intricately orchestrates a myriad of cellular processes pivotal to normal physiological functioning, spanning from cell motility and proliferation to apoptosis and gene transcription.^2^ The intricate equilibrium of Ca^2+^ homeostasis is perturbed in a variety of cancer types and models, suggesting its essential role in oncogenic transformation and metastasis.^3^ The orchestration of calcium signals is facilitated by a wide repertoire of channels, pumps, and exchangers, many of which show notable overexpression in particular cancer phenotypes.^2 3 4^ The upheaval of the Ca^2+^ signaling system is a critical factor that makes cancer cells resistant to treatment interventions, including new molecular and immune-based strategies.^3^ The resistance of chemotherapeutic drugs has been found to be significantly impacted by elevated Ca^2+^ levels and a modified state of Ca^2+^ homeostasis.^4^ Given this, several Ca^2+^ release-activated Ca^2+^ (CRAC) transporter blockers, including carboxyamidotrizol and CM-4620, are presently undergoing clinical studies, either alone or in combination with other therapies, to sensitize tumors to treatment.^5^

Store-operated Ca^2+^ entry (SOCE) stands as an omnipresent and indispensable pathway governing the delicate equilibrium of intracellular Ca^2+^ levels.^6^ This intricate mechanism of calcium influx finds its canonical mediation through ORAI1, a calcium channel ensconced within the plasma membrane, with its activity finely modulated by the dynamic calcium reservoirs of the endoplasmic reticulum (ER). ^6 7 8^ Upon depletion of ER calcium stores, a cascade is triggered wherein the ER Ca^2+^ sensor, stromal interacting molecule 1 (STIM1), undergoes oligomerization and translocation to specialized ER-plasma membrane junctions. Here, at these pivotal cellular crossroads, STIM1 engages with ORAI1, orchestrating a localized surge of calcium influx that underpins myriad cellular processes.^6 9^ The proliferation and migration of several basal breast cancer cells are well-regulated by ORAI1. Interestingly, the heightened expression of the ORAI1 calcium channel emerges as a hallmark feature within the context of triple-negative breast cancer (TNBC), underscoring its indispensable role in fueling the migratory potential and metastatic dissemination of malignant breast tumor cells. Indeed, targeting ORAI1 holds significant promise as a novel therapeutic strategy for combating cancer metastasis and overcoming chemoresistance.^9^ Research endeavors to clarify the molecular mechanisms of ORAI1-mediated signaling in cancer are ongoing; nonetheless, the transcriptional control of this gene is still mysterious.

Our aim was to explore the vicinity of the transcription start site in the promoter region to understand the transcriptional regulation and detect the existence of druggable targets such as G4, which may be subject to selective regulation in cancer treatment. A putative promoter G4 motif, *ORAI1-Pu*, was identified through bioinformatic analysis of the *ORAI1* promoter. Along the line of discovery, G4 structures are among the most significant non-canonical structures in our genome, which are involved in the regulation of intricate biochemical processes. G4s are formed from guanine-rich repeat motifs; their physicochemical characteristics and structural diversity suggest that they are effective therapeutic targets and are highly selective.^10^ They consist of π-π stacked G-quartets with Hoogsteen hydrogen bonding and are stabilized by cations like Na^+^ and K^+^. ^10 11^ Guanine-rich repeats have been discovered in several genomic areas, including telomeres, ribosomal DNAs, immunoglobulin heavy chain switch regions, microsatellite and minisatellite repeats, and the promoter regions of various oncogenes. G4s have been identified as critical agents in the complex regulation of gene expression, influencing several genes at the transcriptional and translational levels.^10 12 13^ Stabilization of these structures offers a promising new area for the development of innovative anti-cancer treatments.^14 15^ The nexus between these G4 structures and the six primary hallmarks of cancer has already been established.^16^ Meanwhile, metabolic reprogramming has emerged as an exclusive hallmark of cancer in recent years. But as of yet, no G4 has been reported to be in the promoter of a membrane transporter gene, which is crucial for metabolic homeostasis and significant signal transmission during oncogenesis.

Since ORAI1 actively contributes to the promotion of drug resistance in cancer cells, further investigation into the *ORAI1-Pu* G4 motif can emerge as a new therapeutic target for the manipulation of further combination therapies. The *ORAI1-Pu* G4 sequence had a propensity to form a G4 structure with parallel topology and an uncommon dynamic 8-nt loop, according to an extensive biophysical characterization and structural determination. Additional research on ligand interaction and in-cellular investigations revealed the endogenous existence of the G4, which, upon probable ligand induced stabilization, reduced promoter activity. Due to the presence of the E-box motif, the dynamic loop region may also be a likely docking site for different transcription factors, such as Zeb1,^17^ c-Myc,^18^ etc. Chromatin Immunoprecipitation (ChIP) data validated the recruitment of Zeb1 in the *ORAI1-Pu* G4 region. It has been found that *Zeb1* is a highly expressed oncogene and facilitator of EMT in TNBC, and the upregulation of *ORAI1* might be a function of binding to E-box presented as a docking site at the *ORAI1-Pu* G4 region. Finally, confocal analysis provided additional support for our hypothesis regarding the functional consequences of stabilizing *ORAI1-Pu* with G4 stabilizing agents, TMPyP4 and BRACO-19. Fura2 detection of calcium uptake also revealed a decrease in SOCE activity in both MDA-MB-231 and B16 cells, indicating that this effect is not limited to a specific species. Overall, our study reports the first-ever proof-of-concept therapeutic module of G4-based anticancer therapeutics targeting Ca^2+^ dynamics.

## Materials and Methods

### Bioinformatics Analysis

The promoter region of the *ORAI1* gene sequence (NCBI database Gene ID: 84876) was analyzed and endorsed by the online Eukaryotic Promoter Database (EPD).^19^ The putative G4 motifs were identified, and their potential score to form a quadruplex was calculated using the QGRS G4 predictor software tool.^20^ Further, the Encode database and UCSF browser were studied to analyze the ChIP-Seq elements near the putative G4 sequence *ORAI1-Pu* with respect to the Transcription Start Site (TSS).^21 22^ The EPD transcription factor motif search tool used the JASPER CORE database 2018^23^ to bioinformatically predict the transcription factors with putative interaction at the *ORAI1-Pu* G4 motif (p-value cutoff set at < 0.01). The 100 vertebrates’ conservation by PhastCons Alignment tool in the UCSF browser was used to study conversed regions within the predicted sequence.

### Sample preparation

All DNA oligonucleotides for structure calculations were purchased from TIB MOLBIOL (Berlin, Germany) and purified by precipitation with potassium acetate and ethanol. The concentration of DNA oligonucleotides was determined by measuring the UV absorbance in water at 260 nm at 80 °C. All DNA oligonucleotides were dissolved in 10 mM potassium phosphate buffer, pH 7. Samples were annealed by heating to 90°C for 5 minutes, followed by slow cooling to room temperature. Final sample concentrations were 5 μM for UV and CD measurements and ranged from 0.3 to 0.4 mM for the NMR studies.

The cationic porphyrin derivative 5,10,15,20-tetrakis (1-methyl-4-pyridinio) porphyrin tetra (p-toluenesulfonate) (TMPyP4) was obtained from Sigma-Aldrich (catalog no. 613560). The synthetic acridine analogue BRACO-19 was obtained from Sigma-Aldrich (catalog no. SML0560) and dissolved in ultrapure water (Invitrogen) at a high stock concentration of 20 mM and stored in the dark at 20 °C.

### Thermal stability determined by UV melting experiments

UV melting experiments for the G4s were performed using a Jasco V-650 spectrophotometer (Jasco, Tokyo, Japan) equipped with a Peltier thermostat. The absorbance of the oligonucleotides (∼5 μM) was recorded at 295 nm from 10 to 90 °C at a heating and cooling rate of 0.2 °C/min and a bandwidth of 1 nm using quartz cuvettes of 1-cm path length. Melting temperatures *T*_m_ were determined from the minimum of the first derivative of the heating curve. All experiments were performed in triplicate.

### Topology prediction by CD spectroscopy

CD spectra were acquired using a Jasco J-810 spectropolarimeter (Jasco, Tokyo, Japan) equipped with a thermoelectrically controlled cell holder. CD spectra were obtained by accumulating five scans recorded at a rate of 50 nm/min over a range of 220-320 nm. The bandwidth was 1 nm and the response time was 4 seconds. Oligonucleotides (∼5 μM) were measured in 1-cm quartz cuvettes at 20 °C in 10 mM potassium phosphate buffer, pH 7. All spectra were blank-corrected by subtraction of the buffer spectrum.

### Structural determination by NMR Spectroscopy

All NMR spectra were acquired on a Bruker Avance NEO 600 MHz NMR spectrometer equipped with an inverse ^1^H/^13^C/^15^N/^19^F quadruple resonance cryoprobe head and z-field gradients. Oligonucleotides (0.3-0.4 mM) were dissolved in 10 mM potassium phosphate buffer, pH 7.0, with 90% H_2_O/10% D_2_O. Spectra were either acquired at 25 °C or 40 °C. Topspin 4.0.7 and CcpNmr Analysis 2.4.2 were used for spectral processing and analysis.^24^ The chemical shift of protons was indirectly referenced to sodium trimethylsilyl propionate (TSP) through the temperature-dependent water chemical shift at pH 7.0. Chemical shifts of carbon were referenced to sodium trimethylsilylpropanesulfonate (DSS) through an indirect referencing method. For details on the NMR experiments and acquisition parameters used see the Supporting Information.

### NMR structure calculations

Initially, 400 structures were calculated using a simulated annealing protocol in XPLOR-NIH 3.0.3 and 100 lowest-energy starting structures were used for further refinement.^25^ The refinement was carried out in vacuum using AMBER18^26^ and was followed by simulated annealing to obtain twenty converged structures. For an additional refinement in water, ten lowest-energy conformations were selected. After neutralizing the system by the addition of potassium ions, two additional potassium ions were placed within the inner channel of the G4 core and the system hydrated with TIP3P water.^27^ The trajectory of a final simulation of 4 ns at 1 atm and 300 K was averaged over the last 500 ps and minimized in vacuum to obtain the final ten lowest-energy structures. The calculated structures were analyzed by using the VMD 1.9.2 software. Pymol 1.8.4 was used for the three-dimensional structural representations. For a detailed protocol of the simulations including settings for the NMR restraints see the Supporting Information.

### Mammalian Cell culture

MDA-MB-231 and A549 cell lines were grown in Dulbecco’s Modified Eagle Medium (DMEM) media (Gibco Catalog No. 11995-065) supplemented with 10% (v/v) FBS (Gibco Cat. No.-10082147), antibiotics Gentamicin (75 μg/ml), 1% Pen-Strep, and Amphotericin B (0.375 μg/ml). They were maintained and passaged in tissue culture treated T-25 flasks and all subsequent experiments done after 2-3 cell passages. Cells were kept under humidified conditions at 37 °C and 5% CO_2_.

### Promoter Activity by Luciferase assay

The upstream promoter region of the human *ORAI1* gene with the putative G4-forming sequence *ORAI1-Pu* was amplified by PCR using human genomic DNA as a template and then cloned into the pGL4.72[hRlucCP] vector (Promega; Madison, USA; catalog no. E6901) at the KpnI/HindIII site. The construct, named *ORAI1*-G4-WT, contained a 300 bp (-259 to +41 from TSS) sequence with the proximally placed *ORAI1-Pu* G4 sequence. Its mutant variant, named *ORAI1*-G4-Null, was synthesized by deleting the *ORAI1-Pu* sequence **(Figure 3A)**. Both the constructs were outsourced from GenScript, USA. For the reporter assay, 0.5 X 10^4^ cells were seeded on a 96-well culture plate. After 24 h, each pGL4.72[hRlucCP] construct (wild-type or mutated, 100 ng) was transiently transfected into cells in Opti-MEM media (HiMedia) with Lipofectamine 2000 (Invitrogen). Each well was also transfected with a pGL3 vector (10 ng, Promega) as a transfection control. After 4 h of transfection, the reduced serum medium used for transfection was replaced with a fresh complete one, and appropriate doses of TMPyP4 or PBS (for control cells) were added to the cells, which were further incubated for the next 24 h. On the following day, cells were harvested and lysed with 1× Passive Lysis buffer (Promega) to determine Firefly and *Renilla* luciferase activities using a Dual-Luciferase Reporter Assay System (Promega) in a Luminoskan luminometer (ThermoScientific). *Renilla* luciferase activity was normalized with Firefly luciferase activity for each sample and considered as relative luciferase activity. The fold change of wildtype was calculated to the G4 null mutant values and the p-value (by paired t-test). Graphical representation was done using GraphPad Prism 8.

### RNA expression analysis

Cells were grown in 6-well plates to 60% confluency, followed by TMPyP4 (25 μM and 50 μM) and BRACO-19 (25 μM and 50 μM) treatment for 24 hours. The cells were lysed and extracted in a Trizol RNA-stabilizing solution. RNA was extracted subsequently using chloroform and isopropanol following a standard protocol. Estimation of RNA was done using Nanodrop. First-strand cDNA synthesis was done with Verso cDNA synthesis kit (Thermo Scientific) followed by a quantitative Real-time PCR assay by using PowerUp SYBR Green Master Mix (Applied Biosystems-Thermo Scientific). As a basal constitutive expression control, the 18S gene expression was quantified. The PCR primers used to quantify *ORAI1* and 18S rRNA are given in **Table S5**. The annealing temperature used was 54°C. Quantification expression was done by the Fold enrichment method; the equations used are:

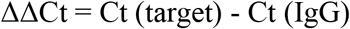

Fold enrichment: 2^^(-ΔΔCt)^

Their statistical significance p-value (by paired t-test) and graphical representation were computed in GraphPad Prism 8.

### Chromatin immunoprecipitation (ChIP)

The chromatin immunoprecipitation assay (ChIP) was conducted to monitor the occupancy level of RNA polymerase and various E-box binding oncogenic transcription factors (Myc22, Zeb1, SNAI2) at the *ORAI1-Pu* promoter region of the *ORAI1* gene promoter.^28^ We also monitored the binding of BG4, which is an anti-G4 antibody, a probable cellulose predictor of G4 formation at *ORAI1-Pu*. A negative control set with an anti-IgG antibody was also performed to eliminate effects due to nonspecific binding. We have used the Thermo Scientific™ Pierce™ Magnetic ChIP kit (Cat. # 26157) to perform the ChIP experiment. MDA-MB-231 cells were seeded at a density of 1 × 10^6^ cells per 100 mm culture dish and harvested after 24 h incubation in a CO_2_ incubator. Cells were processed, and immunoprecipitation was carried forward as per the directed protocol in the Pierce™ Magnetic ChIP kit product sheet. The details of the antibodies used for the co-immunoprecipitation and the concentrations used are given in **Table S6**. We further proceeded to Real-Time qPCR amplification with the purified ChIP genomic fragments. Real-Time qPCR was performed in a QuantStudio 5 thermal cycler to amplify the collected purified DNA using PowerUp™ SYBR™ Green Master Mix (Cat. # A25742) as per the manufacturer’s protocol. Designed PCR primers encompassing the *ORAI1-Pu* sequence were used to amplify the *ORAI1* promoter region (**Figure 4A, Table S5)**. The thermal cycler was set at the following conditions: initial denaturation step by heating at 95 °C for 10 min followed by 45 cycles of 30 s initial denaturation at 95 °C, 30 s at 55 °C annealing temperature, and 30 s of extension at 72 °C.

Further statistical analysis was performed using a paired t-test in GraphPad Prism version 8 software.

### Calcium imaging

Calcium imaging was performed as reported earlier ^29 30^. Cells were cultured on confocal dishes (SPL life sciences) to attain 60-80% confluency. Cells were pre-treated with either 50 μM TMPyP4/BRACO-19 or vehicle control for 48-72 hrs. Cells were then incubated in a culture medium containing 4 μM Fura-2AM for 30 min at 37°C, 5% CO_2_. Post-incubations, cells were washed 3 times and bathed in HEPES-buffered saline solution (2 mM CaCl_2_, 1.13 mM MgCl_2_, 140 mM NaCl, 10 mM D-glucose, 4.7 mM KCl, and 10 mM HEPES, pH 7.4) for 5 min. Further, 3 washes were given and cells were bathed in HEPES-buffered saline solution without 2 mM CaCl_2_ to ensure the removal of extracellular Ca^2+^ before starting the measurements. A digital fluorescence imaging system (Nikon Eclipse Ti2 microscope coupled with a high-speed PCO camera) was used and fluorescence images of several cells were recorded and analyzed. Excitation wavelengths of 340 nm and 380 nm were alternately employed for Fura-2AM and the emission signal was recorded at 510 nm.

## Results

### Identification of a putative G4 motif at the promoter region of *ORAI1*

A putative G4 motif, *ORAI1-Pu*, was predicted by the QGRS mapper (G score: 37) at the promoter region of the human *ORAI1* gene (NCBI database Gene ID: 84876) ∼100 bp upstream of the TSS in the sense strand as denoted by the Eukaryotic Promoter Database (EPD) (Supplementary Fig1). Further, UCSC Genome Browser data on Human (GRCh38/hg38) from chromosome 12: 12626399-122626500, encompassing *ORAI1-Pu* region, which starts at Ch12: 121626426 position gives in-depth role of the putative G4 in regulation of transcription of our gene of interest *ORAI1. ORAI1-Pu*: 5’ **TGGGCGGGGCACAGGTGGGCGGGG** 3’, was seen to be a region of H3K4me3 methylation site as indicated by ChIP-seq data (Barski et al.),^31^ which has role in RNA polymerase II promoter-proximal pause-release,^32^ also this region indicated to be probable binding site to RNA polymerase subunit 1 which has a DNA binding domain. The sequence of our interest, *ORAI1-Pu*, also incorporates the hexanucleotide consensus E-box motif 5’-CANNTG-3’,^33 34^ making it a potential trans element with an effective role in basic helix-loop-helix-like transcription factor binding.^35^ Therefore, *ORAI1-Pu* presented as a putative G4 forming region, with an indicative role as a promoter regulatory element with a loop region containing an E-box. The G4 motif was further seen to be conserved in primates, and except for variations in the loop regions, it was mostly conserved in other mammals, indicating its evolutionary significance in the regulation of the *ORAI1* gene. **(Figure S1)**

### The *ORAI1-Pu* sequence forms a parallel G4 structure

The circular dichroism (CD) spectral signature of the wild-type *ORAI1-Pu* sequence (5’-TGGGCGGGGCACAGGTGGGCGGGG-3’) features negative and positive amplitudes at around 243 and 263 nm, respectively, typical of a parallel G4 with exclusive homopolar tetrad stacking as a result of three connecting propeller loops (**Figure 1A, B**). In addition, its imino proton spectral region shows twelve well-resolved Hoogsteen-type G imino proton resonances between 11.0 and 12.0 ppm, indicating the formation of a single G4 structure with three G-tetrad layers (**Figure 1C top**).

**Figure 1.**
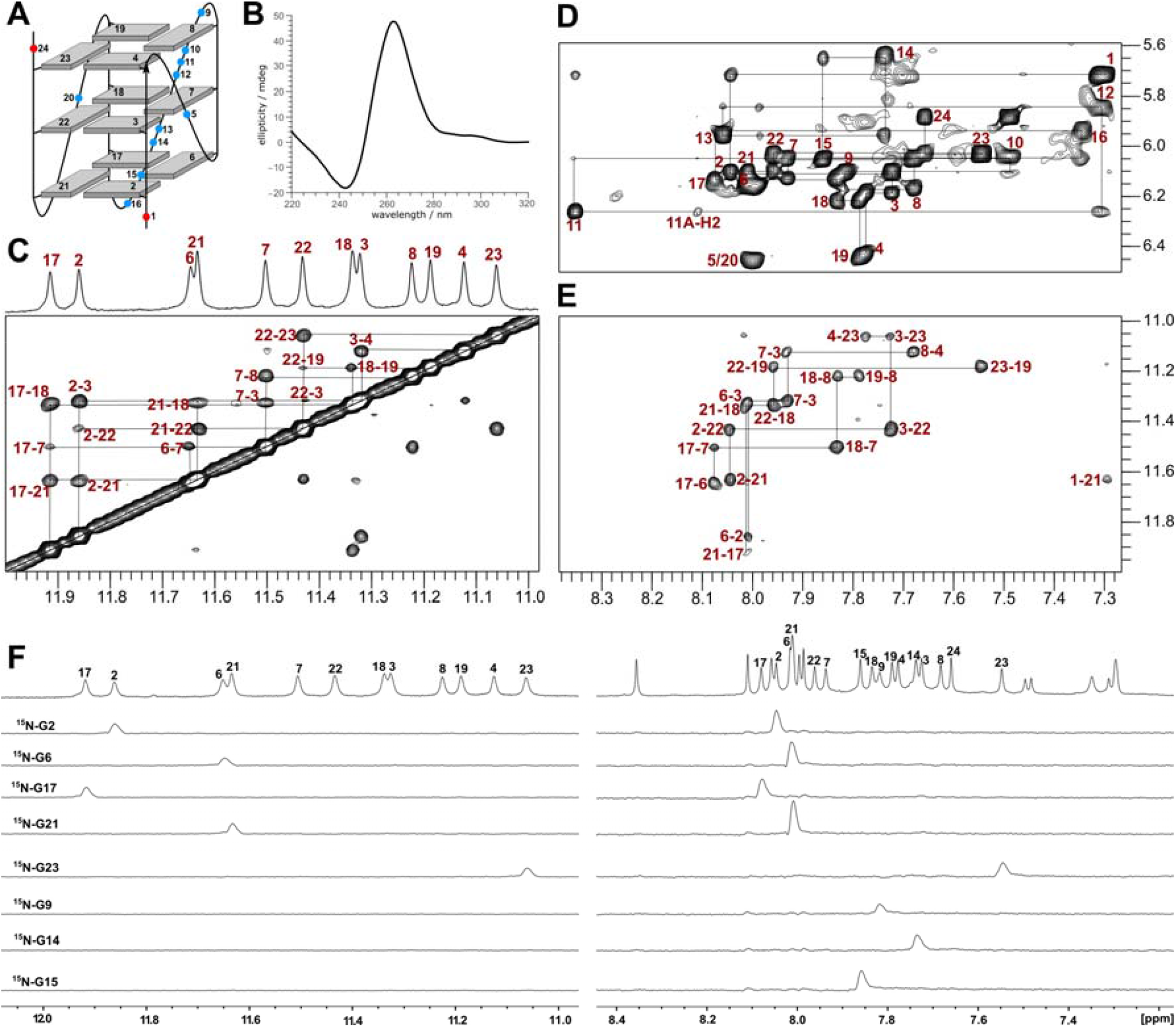
Structure determination of the *ORAI1-Pu* G4. **(A)** Schematic representation of the *ORAI1-Pu* topology with residue numbering; *anti*-G residues of the G-core are colored grey, loop and overhang residues are represented by circles colored in blue and red, respectively. **(B)** CD spectrum of the *ORAI1-Pu* sequence. **(C-E)** 2D NOESY spectral regions of *ORAI1-Pu* (mixing time 300 ms). **(C)** H1-H1 spectral region with corresponding 1D imino proton spectrum shown on top with resonance assignments. **(D)** H8/H6(ω_2_)-H1’(ω_1_) spectral region. **(E)** H8(ω_2_)-H1(ω_1_) spectral region. **(F)** 1D ^15^N-filtered HMQC spectra of *ORAI1-Pu* sequences site-specifically labeled at G2, G6, G17, G21, G23, G9, G14, and G15 (10% ^15^N enrichment). Assignments of guanine H1 (left) and H8 resonances (right) with G imino and G H8 spectral regions of unlabeled *ORAI1-Pu* shown on top. NMR spectra were acquired in a 10 mM potassium phosphate buffer, pH 7.0, at 40 °C.

### NMR structure determination of the *ORAI1-Pu* sequence

A detailed structural characterization of the *ORAI1-Pu* sequence was carried out employing standard NMR methodologies. Full resonance assignments of the folded *ORAI1-Pu* sequence were obtained by the analysis of NOESY, DQF-COSY, ^1^H-^13^C HSQC, and ^1^H-^13^C HMBC experiments. In addition, unambiguous identification of G imino and H8 protons was achieved by site-selective incorporation of ^15^N-labelled G residues **(Figure 1C-F**, for assignment strategies and additional spectra see the **Supplementary Information** and **Figure S2-S4, Table S1)**. NMR-derived distance and torsion angle restraints were used to determine the three-dimensional structure of *ORAI1-Pu* by restrained molecular dynamics calculations in explicit water. **(Table 1)** *ORAI1-Pu* features a parallel topology with a first 1-nt followed by a long 8-nt and another 1-nt propeller loop running in a counter-clockwise direction (**Figure 2**). Such long propeller loops have previously been reported for the parallel G4 in a mutated *c-MYC* and *KRAS* sequence.^36,37^ A superposition of the ten lowest-energy structures shows that the G-core is well-defined (**Table S2**). The resulting structures show significant flexibility along the second 8-nt propeller loop (**Figure S5**). The 5’-terminal residue T1 stacks below G2 and G21 of the 5’-tetrad in agreement with experimental NOE data (**Figure 2C**). 3’-Terminal G24 stacks onto G23 of the 3’-tetrad (**Figure 2D**).

**Table 1:**
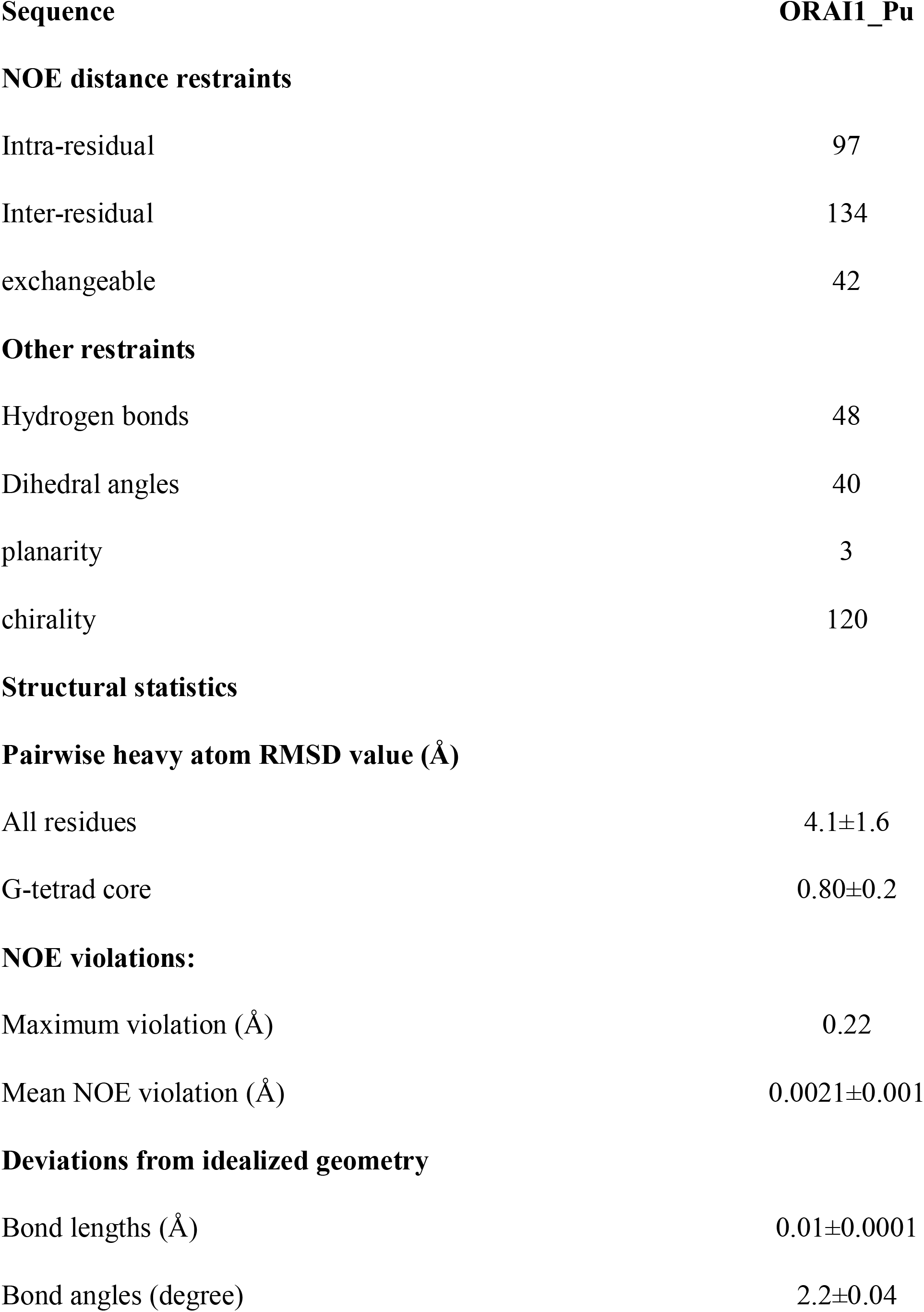
NMR restraints and structural statistics of calculated structures.

| **Sequence** | **ORAI1_Pu** |
| --- | --- |
| **NOE distance restraints** |  |
| Intra-residual | 97 |
| Inter-residual | 134 |
| exchangeable | 42 |
| **Other restraints** |  |
| Hydrogen bonds | 48 |
| Dihedral angles | 40 |
| planarity | 3 |
| chirality | 120 |
| **Structural statistics** |  |
| **Pairwise heavy atom RMSD value (Å)** |  |
| All residues | 4.1±1.6 |
| G-tetrad core | 0.80±0.2 |
| **NOE violations:** |  |
| Maximum violation (Å) | 0.22 |
| Mean NOE violation (Å) | 0.0021±0.001 |
| **Deviations from idealized geometry** |  |
| Bond lengths (Å) | 0.01±0.0001 |
| Bond angles (degree) | 2.2±0.04 |

**Figure 2.**
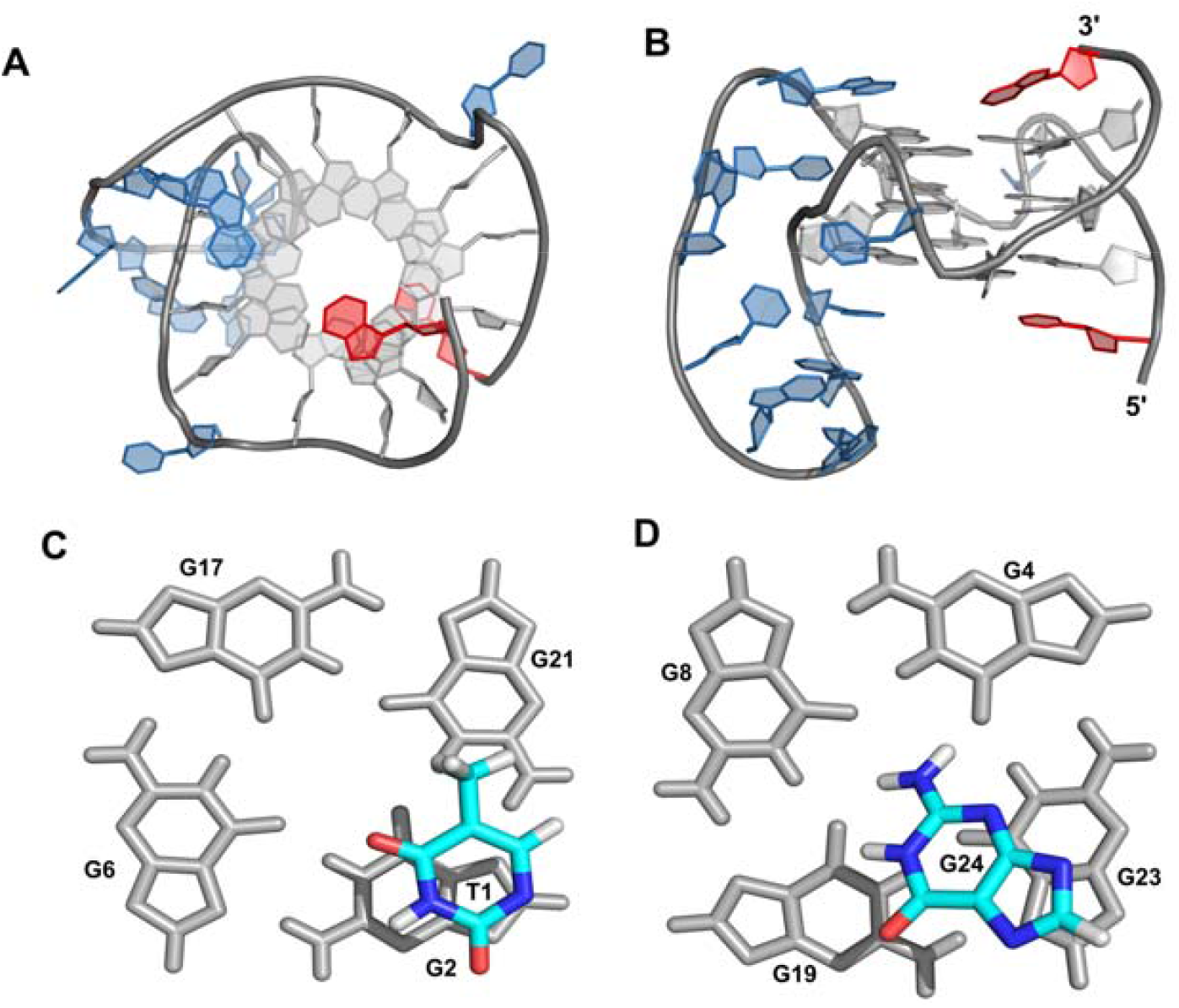
Representative structure of the *ORAI1-Pu* G4. **(A)** Top view and **(B)** side view; *anti*-G residues are colored grey, loop residues are colored blue and overhang residues are colored red. **(C)** View onto the 5’-tetrad with stacked T1 and **(D)** view onto the 3’-tetrad with stacked G24.

### Impact of mutations on the stability of the *ORAI1-Pu* structure

*ORAI1-Pu* mutants were subjected to a CD and NMR spectral analysis **(Figure S6 and S7)**. All mutated sequences exhibit a signature typical of a parallel fold. Additionally, UV melting experiments were performed on all sequences to assess the effect of substituted nucleobases on the stability of the G4 structures. Initially, each residue within the 8-nt propeller loop containing an E-box motif has individually been replaced by a thymidine (**Table S3**). Melting temperatures for all these *ORAI1* single mutants were found to be ∼61 ºC and thus comparable to the melting temperature of the wild type *ORAI1-Pu* sequence determined to be 61.4°C. This demonstrates that a single mutation in the long propeller loop does not affect the stability of the *ORAI1-Pu* structure and suggests the absence of any critical additional interactions of particular loop residues. With all bases in the long propeller loop replaced by thymine in the sequence *ORAI1*_T8, the melting temperature more noticeably decreased by 5 ºC when compared to the wild-type sequence. This can be attributed to reduced short-lived stacking interactions within the long loop when substituting purine by pyrimidine bases. With 3’-terminal G24 replaced by a thymidine in the *ORAI1*_G24T sequence, the melting temperature decreased by 3 ºC. Such a destabilization can be attributed to the favorable stacking of G24 onto the 3’-tetrad in wild-type *ORAI1* as revealed by its high-resolution structure (**Figure 2D**). This indicates that G24 plays a more significant role in the stability of the structure. An additive destabilization with a decrease in melting temperature by 8.3 ºC was found for the *ORAI1*_T8_G24T sequence, exclusively featuring T residues within the long propeller loop as well as a 3’-terminal G24-to-T replacement. Likewise, single and triple G-to-T substitutions in the long propeller loop in addition to a G24-to-T replacement in *ORAI1*_G9T_G24T and *ORAI1*_G9G14G15G24T sequences resulted in a decreased melting temperature by 3.8 ºC and 6.6 ºC, respectively. Taken together, 3’ flanking G24 contributes to the stability of the *ORAI1-Pu* quadruplex by stacking interactions with the outer tetrad, whereas the 8-nt sequence motif in the *ORAI1-Pu* propeller loop results in only rather unspecific and modest stabilizing effects when compared to an all-thymidine loop with the same length.

### Binding of *ORAI1-Pu* motif with various G4 binding ligands-TMPyP4 and BRACO-19

TMPyP4 and BRACO-19 are some common, widely used ligands that show high affinity to bind to G4-like structures.^38,39^ In fact, both TMPyP4 and BRACO-19 show strong binding in the sub-micromolar range to the *ORAI1-Pu* motif and K_d_ values of about 0.1 μM and 0.6 μM were determined by isothermal titration calorimetry **(Figure S8 and Table S4)**. Strong binding of BRACO-19 is also reflected in the increased melting temperature (80 ± 2 °C) upon its addition to *ORAI1-Pu* as observed from the temperature dependent ellipticity at 263 nm **(Figure S9)**. In addition, stacking of BRACO-19 to an outer tetrad of the G-quadruplex is suggested by the appearance of new, albeit broadened, imino signals upon ligand addition that are upfield-shifted compared to the imino resonances of the free quadruplex (**Figure S9**). For TMPyP4, binding seems to follow a more complex behavior possibly comprising various binding modes. Thus, a very broad CD melting transition with an increased temperature (70.4 ± 2 °C) for half-dissociation is observed for a TMPyP4 complex. On the other hand, NMR titrations with TMPyP4 show a gradual loss of free quadruplex imino resonances and the appearance of considerably upfield-shifted signals. Such a behavior may indicate exchange between different ligand binding sites but some minor but also major structural rearrangements of the quadruplex cannot be excluded.

### Effects of the *ORAI1-Pu* motif on promoter activity

The effect of ligand binding on the promoter activity was studied by performing a Dual-Luciferase Reporter assay with a wildtype (with *ORAI1-Pu* motif) and a control mutant (without *ORAI1-Pu* motif) construct. Increasing concentration of TMPyP4 treatment post-transfection in the wildtype construct showed an initial non-significant change at lower concentrations but at higher concentrations ∼30% loss in normalized reporter expression with respect to untreated control. However, for the G4 null mutant construct, at low treatment concentration of ∼5 mM, though there is an initial slight decrease in activity, at further increasing concentrations, there are non-significant changes in reporter activity. In biophysical studies, we have observed that TMPyP4 has a stabilizing effect on the *ORAI1-Pu* structure. Conceivably, TMPyP4 stabilizes the promoter G4 element, i.e., *ORAI1-Pu*, and decreases the promoter activity. The absence of the G4 promoter element in the mutated construct, i.e., G4-null, and the non-significant change in the reporter activity indicate that TMPyP4 does not reduce promoter activity observed by interacting with other probable promoter elements within the -300 bp promoter sequence cloned in the pGL4.72[hRlucCP] vector **(Figure 3B)**. This transcription regulation was further examined in cancer cell-line MDA-MB-231 by quantification of *ORAI1* mRNA level by q-PCR assay upon treatment with both G4 binding ligand TMPyP4 and BRACO-19. The results corroborated with the promoter activity studies done by Luciferase assay as treatment with TMPyP4 and BRACO-19 both decreased expression of the *ORAI1* gene. 25 μM TMPyP4 and BRACO-19 treatment reduced *ORAI1* expression by ∼55% and ∼45%, respectively, which was further highly reduced by greater than 90% (p < 0.05; Fig.) at 50 μM **(Figure 3C)**.

**Figure 3:**
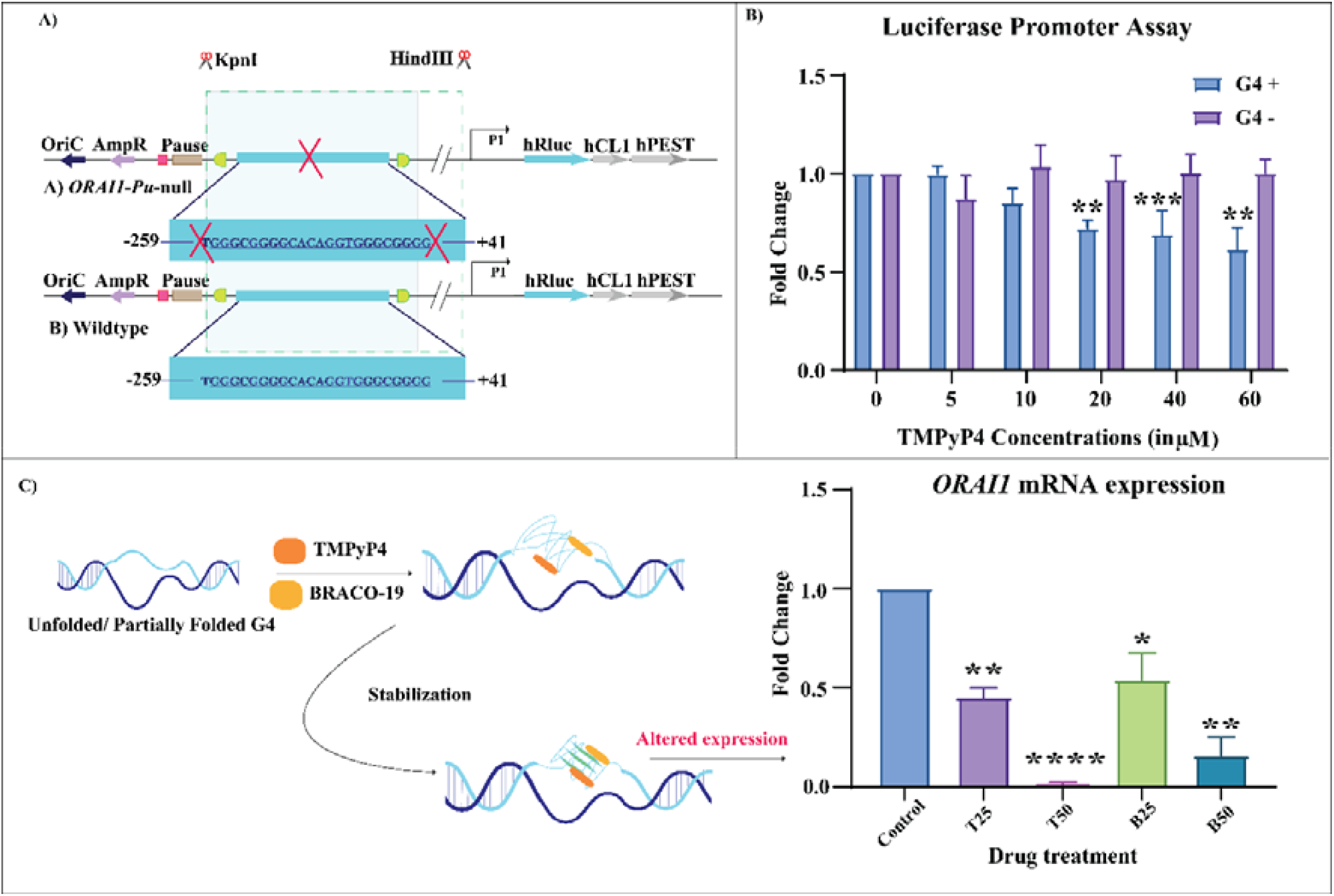
*ORAI1-Pu* G4 as a potential promoter element. **(A)** The representative schematic diagram for the reporter luciferase construct. The promoter sequence of *ORAI1* (249 bp upstream and 41 bp downstream sequence of *ORAI1* from TSS) was cloned with or without the wild-type G4 (*ORAI1-Pu*) scaffold into the Kpn-I and Hind-III restriction sites. Kpn-I and Hind-III restriction sites are upstream of the *hRluc* gene, and the sequence is cloned at the promoter region involved in the expression of the reporter gene *hRluc*. The deletion clone without the *ORAI1-Pu* sequence is called *ORAI1*-*Pu*-null. Abbreviations include ampicillin resistance gene (AmpR), Pause (RNA pol pause signal), oriC (origin of replication), *hCL1* and *hPEST* (protein-destabilizing sequences), and *hRluc* (*Renilla* luciferase gene). **(B)** In MDA-MB-231, the dual luciferase assay was run in increasing concentrations of TMPyP4 using the Wildtype and *ORAI1-Pu*-Null constructs. After comparing the activity of firefly luciferase to that of *Renilla* luciferase, the relative luciferase activity of every clone was computed and displayed as a fold change. **(C)** Representation of effects of G4 ligand. There is stabilization of *ORAI1-Pu*. q-PCR of mRNA post-treatment and quantitative analysis by ΔΔCt shows a loss in *ORAI1* expression on TMPyP4 (25μM & 50μM) or BRACO-19 (25μM & 50μM) treatment. (* represents p-value < 0.05, ** is p-value <0.01, *** is p-value <0.001, **** is p-value <0.0001).

**Figure 4:**
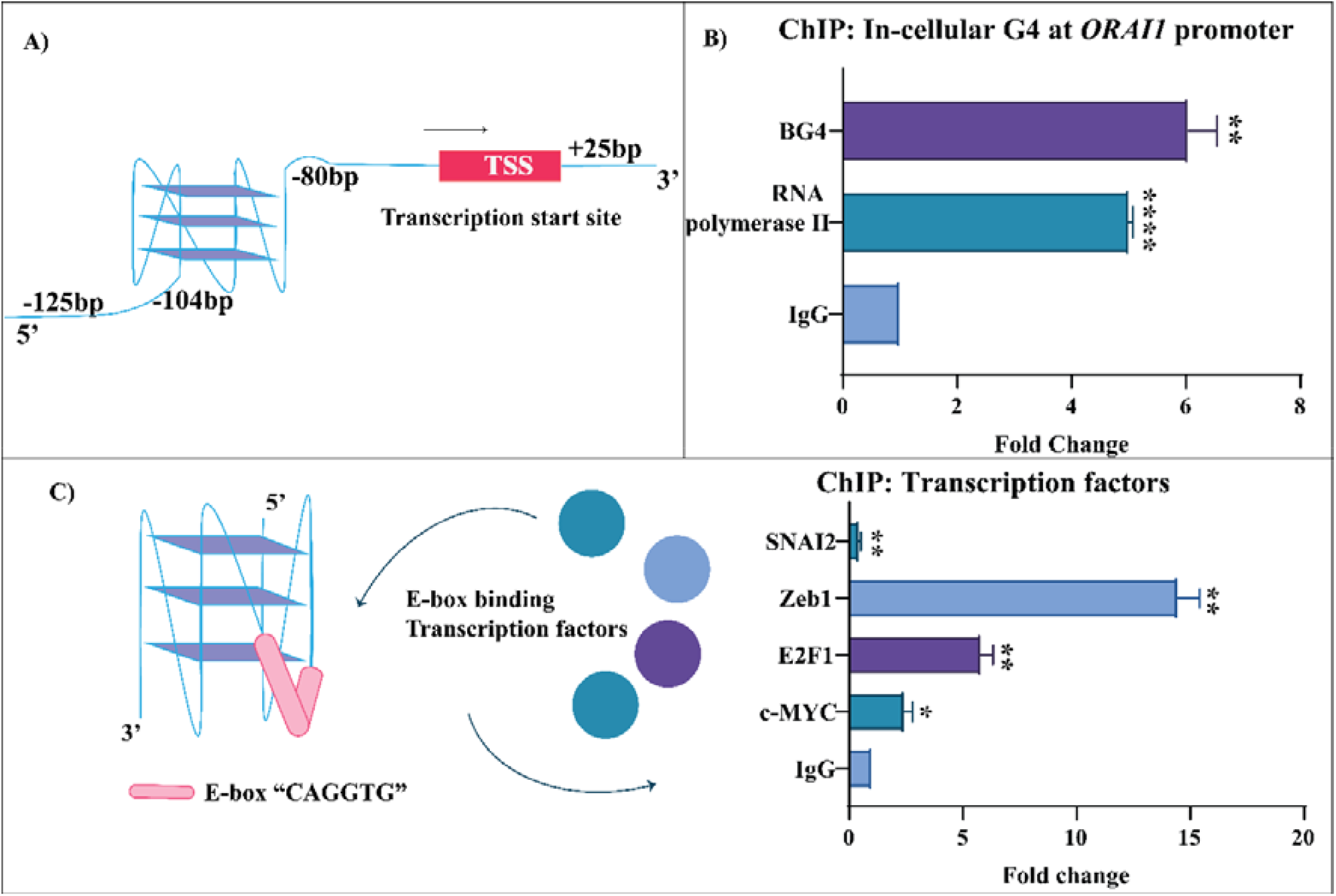
Transcriptional landscape of *ORAI1* promoter mediated by G4. **(A)** Primer design region for ChIP assay encompassing the *ORAI1-Pu* sequence at the *ORAI1* promoter. (B) Endogenous characterization of G4 formation by BG4 at *ORAI1-Pu* and RNA polymerase II binding affinity by subsequent fold enrichment on Chromatin pull-down assay, with IgG as negative control. **(C)** Representation of the region with E-box motif. Transcriptional regulation of *ORAI1-Pu* by acting as a docking platform for various transcription factors (SNAI2, Zeb1, E2F1, and c-Myc) as observed by fold enrichment in ChIP assay. IgG was used as a negative control. (* represents p-value < 0.05, ** is p-value <0.01, *** is p-value <0.001, **** is p-value <0.0001).

### Endogenous study of the *ORAI1-Pu* motif and its probable role as CRE (Cis Regulatory Element)

In order to measure the frequency of the recently discovered G4 formation in chromatin within the cell, we employed the chromatin immunoprecipitation assay (ChIP) with the well-characterized G4 structure-specific antibody (BG4).^41 42^ The PCR primer used encompassed the near promoter region -125 bp upstream to 25 bp downstream of the TSS, which includes our sequence of interest *ORAI1-Pu* **(Figure 4A)**. Pull-down of the region with BG4, compared to IgG, showed significant fold change, indicating the formation of endogenous G4 at the near promoter site of the *ORAI1* gene **(Figure 4B)**. This region also showed a positive interaction with RNA polymerase II on pull-down with RNA polymerase II antibody, evidencing its active role in transcription. The presence of a well-characterized E-box motif 5’-CAGGTG-3’ within the sequence instigated the investigation of endogenous binding of E-box binding transcription factors at this element. Among the transcription factors (Slug, Myc22, Zeb1) studied, Zeb1 showed the highest proficiency for interaction **(Figure 4C)**. We also studied E2F1, which has been characterized to have a major role as a metabolic regulator^43^ and also has a function in the regulation of Zeb1.^44^ *ORAI1-Pu* also encompasses the probable site for E2F1 binding as studied bioinformatically with the EPD transcription factor search tool.^45^ ChIP pull-down assay reveals the plausibility for binding; however, it is lower than that of Zeb1. The results represent that the region has a high likelihood of being a regulatory element by interaction with one or more transcription factors that might be docked onto the non-canonical G4 endogenous structure.

### Loss in SOCE activity on ligand mediated destabilization of *ORAI1-Pu*

To examine the role of G4 binding ligands TMPyP4 and BRACO-19 on *ORAI1* functioning, we performed classical SOCE measurements. *ORAI1* is a well-established store-operated Ca^2+^ entry channel in a variety of cell types, including MDA-MB-231 human breast cancer cells.^9 46^ Therefore, we pre-treated MDA-MB-231 cells with TMPyP4 and BRACO-19 for 72 hrs and performed live cell calcium imaging experiments. Cells were stimulated with 2μM Thapsigargin (Tg) in the absence of extracellular Ca^2+^. Tg inhibits ER localized Sarcoplasmic/Endoplasmic Reticulum Ca^2+^ ATPase (SERCA) pump, thereby releasing ER Ca^2+^ stores. Then, we added 1 mM CaCl_2_ in the bath solution and this resulted in Ca^2+^ entry via SOCE channels present on the plasma membrane, i.e., *ORAI1*. In the first set of experiments, we treated cells with either BRACO-19 or vehicle control nuclease-free water (NFW) and performed live cell imaging. As evident in the traces, BRACO-19 treatment led to a significant decrease in SOCE in MDA-MB-231 cells **(Figure 5A)**. We repeated the experiment in multiple independent runs and the data from hundreds of cells showed an about 50% reduction in *ORAI1*-mediated SOCE **(Figure 5B)**. Next, we performed similar imaging experiments with pre-treatment of TMPyP4. We observed the same trend upon treatment with TMPyP4, i.e., SOCE is clearly decreased upon TMPyP4 treatment **(Figure 5C)**. After performing multiple independent runs, the data from over three hundred cells suggest over 40% abrogation of SOCE **(Figure 5D)**. Taken together, these data clearly demonstrate that modulation of G4 with TMPyP4 and BRACO-19 results in a substantial reduction in *ORAI1* functioning in MDA-MB-231.

**Figure 5:**
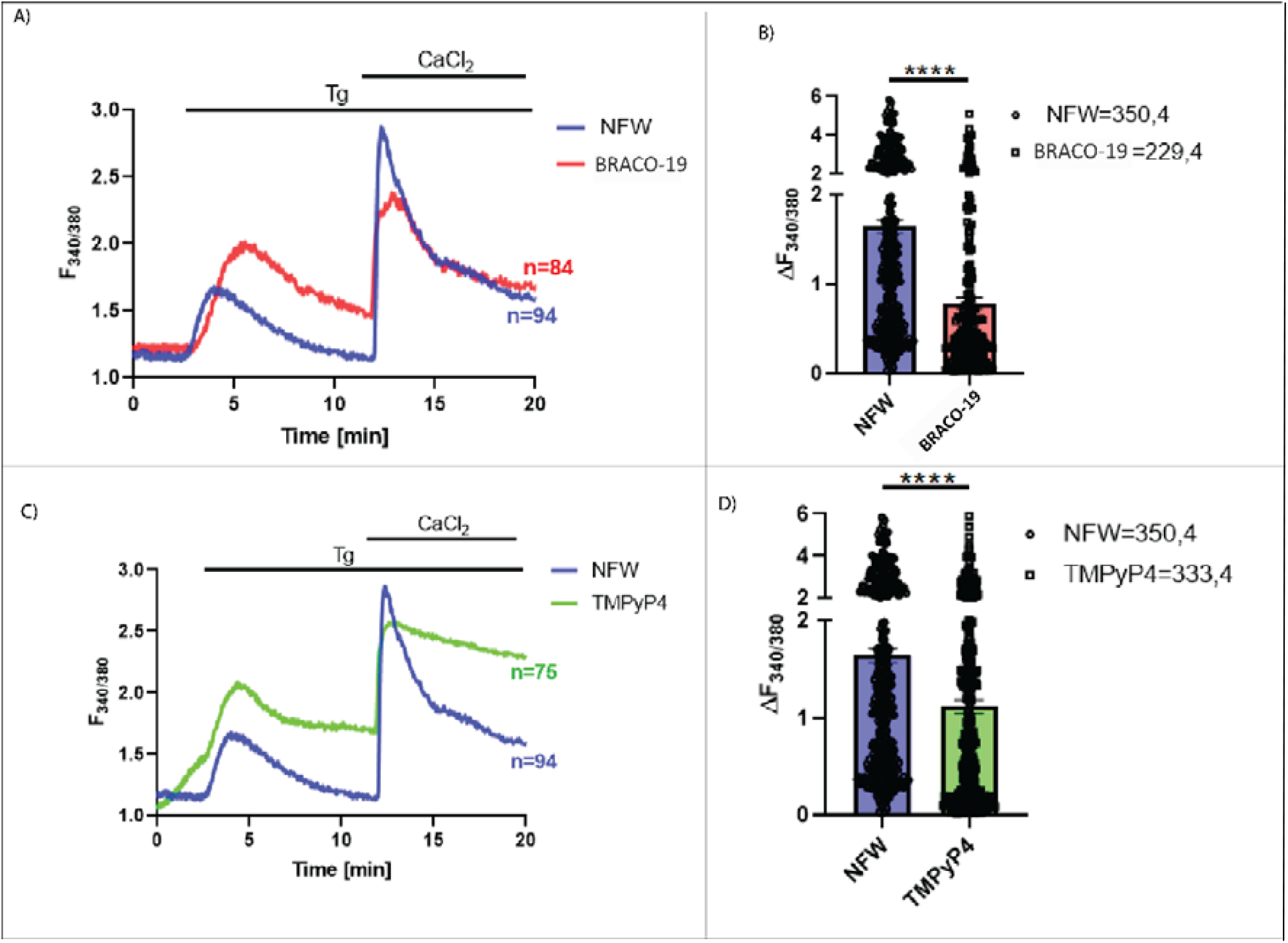
Downregulation of SOCE activity by G4 binding ligands at *ORAI1*-*Pu*. **(A)** and **(C)** Representative Ca^2+^ imaging traces for control Nuclease Free Water i.e. NFW (n=94) and treatment **(A)** BRACO-19 (n=84) or **(C)** TMPyP4 (n=75) where ‘‘n’’ denotes the no. of cells in that trace. Cells were stimulated with 2 μM thapsigargin (Tg) in Ca^2+^ free buffer and restored with 1 mM extracellular Ca^2+^. In **(B)** and **(D)**, the extent of SOCE was calculated from 350 NFW and **(B)** 229 BRACO-19 or **(D)** 333 TMPyP4 treated MDA-MB-231 cells, which were imaged from 4 independent experiments (“n = x, y” where “x” denotes a total number of cells imaged and “y” denotes the number of traces recorded). Data presented are mean ± S.E.M. For statistical analysis, an unpaired student’s t-test was performed. Here, **** means p< 0.0001.

To further substantiate the effect of G4 modulation on *ORAI1* functioning, we measured SOCE in B16 mouse melanoma cells. We have already reported that *ORAI1* constitutes a functional SOCE channel in these cells.^47 48^ We pre-treated B16 cells with TMPyP4 and carried out live cell Ca^2+^ imaging at 48 and 72 hrs post-treatment. As observed in MDA-MB-231 cells, TMPyP4 treatment results in a significant decrease in *ORAI1*-mediated SOCE in B16 cells as well **(Figure S10A)**. The quantitation of SOCE from multiple independent runs in over two hundred cells demonstrates that TMPyP4 treatment time-dependently reduces SOCE in B16 cells with above 40% decrease at 48 hrs and around 60% abrogation at 72 hrs **(Figure S10B)**. Collectively, these data elegantly corroborate that the G4 modulation results in a robust decrease in *ORAI1* function in B16 cells. Importantly, data from human MDA-MB-231 cells and mouse B16 cells reveal that the effect of G4 modulation on *ORAI1* function may be similar in various cell types and species. It is rather a conserved phenomenon observed in cell lines originating from different tissues and species.

Being another interesting observation from the Ca^2+^ imaging experiments, the treatment with TMPyP4 and BRACO-19 also results in higher ER Ca^2+^ release, which does not come back to baseline as seen in the control NFW condition. This suggests that apart from *ORAI1*, TMPyP4 and BRACO-19 may also be acting on G4 close to other Ca^2+^ handling proteins such as Plasma Membrane Ca^2+^ ATPase (PMCA) and ER Ca^2+^ leak channels. However, future studies would be required to decipher the comprehensive effect of G4 modulators on cellular Ca^2+^ signaling via their action on other channels/transporters.

## Discussion

In our study, we found evidence that the *ORAI1* gene promoter had the propensity for the formation of regulatory non-canonical G4 DNA structures. The G4 forming motif was evolutionarily conserved and shows twelve well-resolved Hoogsteen-type G imino proton resonances between 11.0 and 12.0 ppm, indicating the formation of a single G4 species with three G-tetrad layers. Determination of the high-resolution G4 structure revealed that it constitutes a parallel-stranded G4 with three loops and exclusive homopolar tetrad stacking. The first loop was 1-nt long, followed by a long 8-nt loop and another 1-nt propeller loop running in a counter-clockwise direction. The presence of the long 8-nt loop is unusual in parallel G4s **(Figure 2)**. However, this region is a highly dynamic single-stranded region and has an E-box-like motif 3’-CAGGTG’ that might be a probable binding site for transcription factors **(Figure S5)**.^49^ Mutation studies with this sequence revealed that single loop nucleotide mutations did not change the *T*_m_ of the G4 significantly, suggesting the absence of any critical additional interactions at particular loop residues for the stability of the structure **(Table S3)**. TMPyP4 and BRACO-19, two commonly used G4 binders, were used to study the effects of stabilization of the G4 on promoter activity. Both the ligands showed an affinity to bind to the putative G4 and stabilized it, as seen by an increase in thermal stability by CD spectroscopic melting experiments. **(Figure S8 and S9)** It was found by Dual-Luciferase reporter assay that *ORAI1-Pu* acts as a transcriptional repressor and treatment with the stabilizing ligand TMPyP4 reinforces its regulatory activity. The treatment of these ligands in breast cancer cell line MDA-MB-231 further corroborated our finding as treatment and possible stabilization of *ORAI1-Pu* significantly decreased the mRNA levels of the *ORAI1* gene, which is commonly overexpressed in TNBC cancer subtypes **(Figure 3)**.^7^ However, the exact effects of structural dynamics occurring at this locus are ambiguous and cannot clarify which conformer of the unfolded or partially folded structure regulates transcription. The endogenous presence of the G4 structure was noted by ChIP with widely used anti-G4 antibody BG4 as there was significant interaction in the pull-down region, which was amplified with a primer specific to the *ORAI1-Pu* region. Further ChIP investigation revealed that the region was an RNA polymerase II binding site and possibly provided a docking platform for various transcription factors. The highest propensity of binding was with the E-box binding transcription factor Zeb1 **(Figure 4)**. Zeb1 has been recognized to be an oncogene over-expressed in most cancer cell lines, including MDA-MB-231. It has been associated with regulating various genes to initiate EMT in cancer cells. *ORAI1* is a metabolic Ca^2+^ signaling regulator and has a major role in cancer homeostasis. Zeb1 docking to the *ORAI1-Pu* region might give us new insight into the complex regulation of these oncogenic factors. Since there is higher fold binding of Zeb1 compared to BG4 in MDA-MB-231, the G4 structure is probably partially destabilized resulting in overexpression of *ORAI1*. However, further experiments need to be carried out to understand the partial folding dynamics of the G4 at this site. *ORAI1* mRNA deregulation by TMPyP4/BRACO-19 ligands cascaded on the effects downstream to the depletion of Store Operated Calcium Entry by ∼50% **(Figure 5)**. This observation was evident in both data from human MDA-MB-231 cells and mouse B16 cells **(Figure S10)**, suggesting that the effect of G4 modulation on *ORAI1* function maybe similar in various cell types or species. As the *ORAI1-Pu* sequence is conserved in primates and nearly conserved in mammals, it might be a conserved phenomenon observed in cell lines originating from different tissues and species.

## Conclusion

*ORAI1* is a cellular membrane transporter with a wide array of cellular functions as an important member of the CRAC channel.^8^ The calcium homeostasis that is maintained by *ORAI1* is in a very integrated complex signaling cascade involving a lot of other cellular effectors.^50^ *ORAI1* might mediate its function through store-dependent or independent pathways by interaction with other factors overexpressed in cancer, like SPCA2.^51^ However, though much has been extensively studied about the interaction pathways between the proteome and its subsequent role in cancer,^7^ the transcription regulation of *ORAI1* is still not extensively characterized. The presence of the *ORAI1-Pu* motif and the propensity of regulation of the gene through this element directs us to some insights into the cis-regulatory element of *ORAI1* **(Figure 6)**. The structure of the non–canonical G4 at *ORAI1* promoter can further be studied and presented as a druggable target to regulate the transcription of *ORAI1*.^15^ In our study, we have elucidated the role of G4 using one cancer sub-phenotype of MDA-MB-231. G4s are transient structures, and stabilization/destabilization might be regulated temporally within a cell and spatially within various tissues and subtypes depending on physiological factors.^12^ Therefore, further study of the G4 present at the regulatory element of *ORAI1* can provide us with insight into the regulation pattern of this transporter gene in various alternate signaling phenotypes. Eventually, our study marks a significant milestone by introducing the pioneering proof-of-concept therapeutic framework of G4-based anticancer treatments, specifically tailored to modulate calcium dynamics.

**Figure 6:**
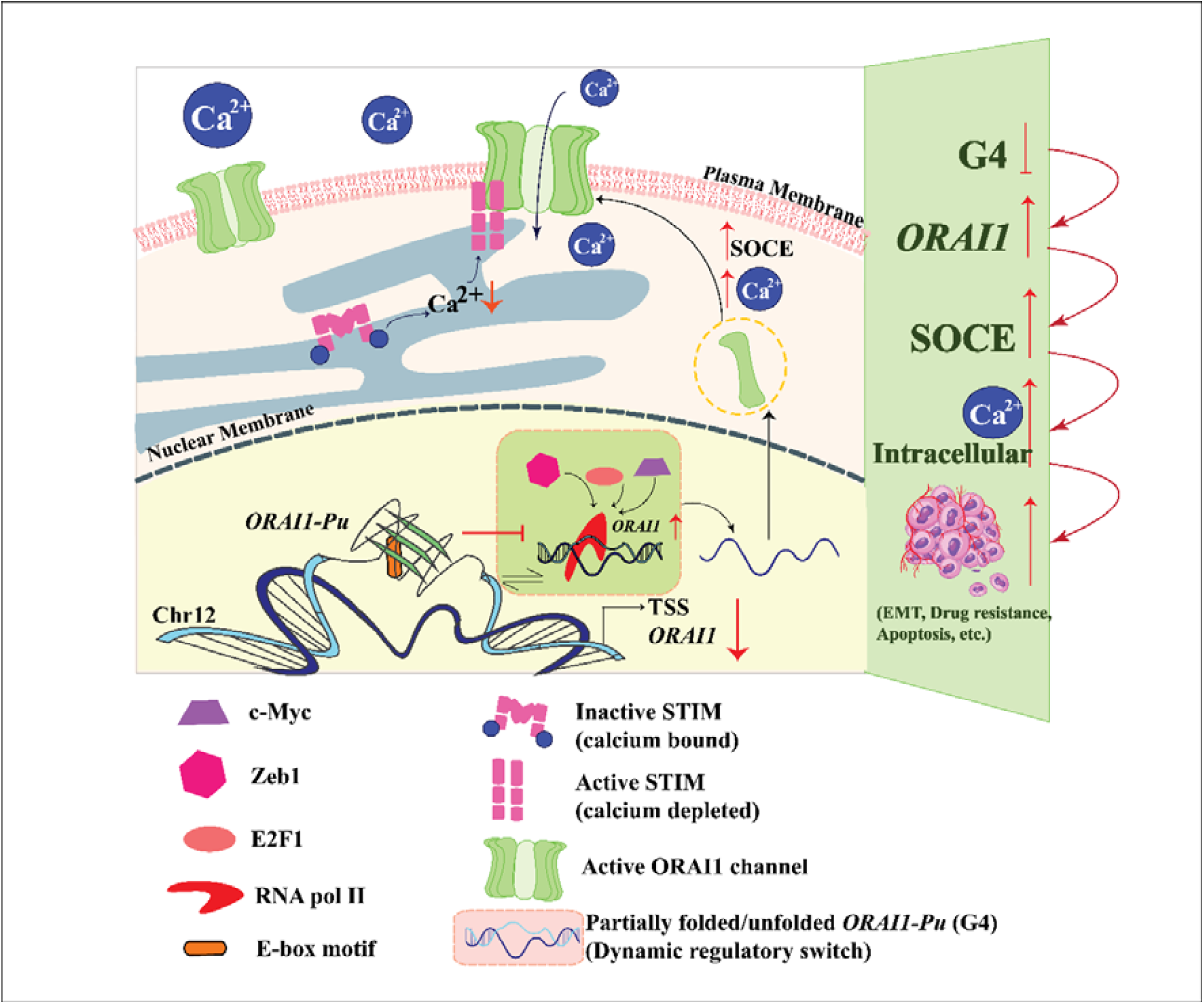
Overview/TOC. *ORAI1* expression is downregulated in the presence of a stable G4 structure at the predicted *ORAI1-Pu* motif, which is formed at a regulatory docking site for transcription factors and RNA polymerase II. The G4 structure might act as a dynamic switch, as it is a transient structure and is regulated by various factors within a cell. Ligand-mediated stabilization of ORAI1-Pu via TMPyP4/ BRACO-19 indicated gain-of-function in *ORAI1* expression. In MDA-MB-231, we observed greater fold endogenous binding of Zeb1 at the ORAI-Pu motif, compared to BG4, which binds to stable G4 structures. In MDA-MB-231, the *ORAI1-Pu* locus might be in a partially unfolded state or unfolded state, leading to over-expression of *ORAI1*. However, further studies can reveal the physiological dynamics of the structure and its functional effects on *ORAI* expression. Overexpression of ORAI1 increases its transport to the plasma membrane and increases SOCE on interaction with activated STIM at physiological conditions where there is loss of Ca^2+^ in the endoplasmic reticulum (ER) calcium store. A spike in intracellular Ca^2+^ cascades its effect on deregulated cellular activity and enhances various hallmarks of cancer, epithelial-mesenchymal transition (EMT), drug resistance, and immune evasion. Etc.

## Supplementary data

Supplementary Data are available at NAR Online.

## Acknowledgements

Central Instrumental Facility (CIF) of Bose Institute and NMR and CIF facility at Universitat Greifswald is highly acknowledged.

## Author contributions

O.C., and S.P.: Conceptualization, Writing – original draft, Writing - review & editing; O.C.: Methodology, In-cellulo and interaction studies, Formal analysis, Software, Visualization; J.J.: Methodology, Structure calculations, Formal analysis, Software, Visualization, Writing – original draft, Writing - review & editing; A.D.: Methodology, Formal analysis; A.S., S.S. and R.M.: Methodology, Calcium imaging, Formal analysis, Writing – original draft; K.W. and S.C.: Supervision, Funding acquisition, Validation, Resources and Project administration, Formal analysis, Writing - review & editing. All authors have given approval.

To the final version of the manuscript. O.C., J.J. and S.P. contributed equally to this work.

