## Supplementary Data for "Remodeling Ca^2+^ dynamics by targeting a promising E-box containing G-quadruplex at *ORAI1* promoter in triple-negative breast cancer"

### Supplementary Information

### Contents:

1. **Structural Determination of *ORAI1-Pu***
   1.1. **NMR experiments**
   1.2. **NMR spectral analysis and resonance assignments of *ORAI1_Pu***
   1.3. **Protocol for NMR-restrained structure calculations**

2. **Biophysical Analysis of Ligand interaction to *ORAI1-Pu***
   2.1. **Isothermal titration calorimetry**
   2.2. **CD melting experiments**
   2.3. **NMR titration experiments**

3. **Supplementary Figures (S1-S10)**

4. **Supplementary Tables (S1-S5)**

### 1. Structural Determination of *ORAI1-Pu*

#### 1.1. NMR experiments

NMR experiments included $^{1}H$-$^{1}H$ nuclear Overhauser effect (NOESY), $^{1}H$-$^{13}C$ heteronuclear single quantum coherence (HSQC), $^{1}H$-$^{13}C$ heteronuclear multiple bond coherence (HMBC), double quantum-filtered correlation (DQF-COSY), and one-dimensional $^{1}H$-$^{15}N$ heteronuclear multiple quantum (HMQC) experiments for $^{15}N$ editing. For solvent suppression in 1D spectra and 2D NOESY experiments, an optimized WATERGATE with a w5 element was used. NOESY spectra were recorded with 300, 150, and 80 ms mixing times. Phase-sensitive HSQC experiments were performed with a 3-9-19 water suppression scheme, 4K×400 data points, and a spectral width of typically 7.5 kHz in the indirect dimension to accommodate C6/C8/C2 resonances of the nucleobase. HMBC spectra were recorded with a jump-and-return water suppression, 2K×144 data points, and processed with 50% non-uniform sampling (NUS) in the indirect dimension. DQF-COSY spectra were acquired using a 3-9-19 water suppression scheme in 90% $H_2O$/10% $D_2O$ and 4K×500 data points.

$^{15}N$ editing was performed using 1D $^{1}H$-$^{15}N$ HMQC experiments on selectively $^{15}N$-labeled oligonucleotides with 10% $^{15}N$ enrichment. For the detection of guanine H1 imino protons, a selective 90° excitation pulse on the imino resonance was used. This was followed by a standard HMQC pulse sequence with $^{15}N$ decoupling during acquisition. The experiment was optimized for an N-H heteronuclear coupling constant of 90 Hz. A modified HMQC pulse scheme, including a 180° shaped $^{15}N$ pulse for selective refocusing of the additional scalar coupling from H8 to N9 and a 90° shaped $^{15}N$ purge pulse, was used to detect guanine H8 resonances. The experiment was optimized for a N-H heteronuclear coupling constant of 16 Hz.

#### 1.2. NMR spectral analysis and resonance assignments of *ORAI1-Pu*

The wild-type *ORAI1-Pu* sequence shows twelve well-resolved imino proton resonances between 11 and 12 ppm, indicating the formation of a single G4 structure with three G-tetrad layers (Figure 1C). All guanine residues in the sequence can be identified as *anti*-guanosines by their intra-residual H8-H1' NOESY cross-peak intensities and upfield-shifted $^{13}C8$ resonances observed in a $^{1}H$-$^{13}C$ HSQC spectrum (Figure S2). A continuous base-sugar sequential NOE walk in the H6/H8-H1' region can be followed from residue T1 to G4 along the first G-column, i.e., G2-G3-G4 (Figure 1D). Another uninterrupted NOE walk begins at G6 and extends to G19. It identifies the second (G6-G7-G8) and third (G17-G18-G19) G columns. The fourth G-column (G21-G22-G23) can be identified by the uninterrupted NOE connections from G21 to the 3' flanking G24. Sequential connectivities are interrupted by the one-nucleotide C5 and C20 propeller loops.

Guanine imino protons of the G-core were unambiguously assigned through $^{1}H$-$^{13}C$ HMBC spectra, correlating H8 and H1 protons via their long-range couplings to $^{13}C5$ at natural abundance (Figure S3). In addition, assignments for G2, G6, G17, G21, and G23 imino and H8 protons were confirmed by $^{15}N$ experiments using residue-specific $^{15}N$-labeled oligonucleotides with 10% $^{15}N$ enrichment (Figure 1F). Further, G9, G14, and G15 H8 protons were again confirmed by corresponding $^{15}N$ experiments with specifically $^{15}N$-labeled sequences. The tetrad polarities were determined using characteristic H8-H1 intra-tetrad NOE contacts (Figure 1E). Assignments were further confirmed by imino-imino NOE contacts (Figure 1C). These were typical of an all-homopolar stack of G-tetrads. Thus, the direction of the Hoogsteen hydrogen bonds from donor to acceptor within the tetrads points along G2-G6-G17-G21 (5'-tetrad), G3-G7-G18-G22 (central tetrad) and G4-G8-G19-G23 (3'-tetrad). This also reveals a parallel topology with two one-nucleotide propeller loops and one eight-nucleotide propeller loop.

Sugar conformations of the residues were determined from DQF-COSY cross peak intensities after stereospecific assignment of H2'/H2" protons in an 80 ms NOESY experiment (Figure S4).

**1.3. Protocol for NMR-restrained structure calculations**

Initially, 400 structures were obtained by a simulated annealing protocol in XPLOR-NIH 3.0.3, keeping 100 lowest-energy starting structures. Distance restraints for the calculations were set according to the intensities of the cross-peaks in 2D NOESY spectra. For non-exchangeable protons, intensities were categorized as strong (2.9 ± 1.1 Å), medium (4.0 ± 1.2 Å), weak (5.5 ± 1.5 Å), and very weak (6.0 ± 1.5 Å). Distances were set to 5.0 ± 2.0 Å in the case of ambiguous cross-peaks due to signal overlap. For exchangeable protons, distances were assigned according to 2.9 ± 1.1 Å for very strong cross-peaks, 4.0 ± 1.2 Å for strong cross-peaks, 5.0 ± 1.2 Å for weak cross-peaks, 6.0 ± 1.2 Å for very weak cross-peaks and 5.0 ± 2.0 Å for ambiguous cross-peaks. Glycosidic torsion angles were restricted to a range of 170-310° for anti-conformers. For south sugar puckers based on experimentally determined scalar couplings from DQF-COSY cross-peak patterns, the pseudorotation phase angle was set to 144-180°. Planarity and hydrogen bond restraints were employed for bases in each G-tetrad.

The refinement was carried out in vacuum using AMBER18 with the parmbsc force field and OL15 modifications for DNA. Restraint energies were set to 40 kcal·mol$^{-1}$·Å$^{-2}$ for NOE-based distance restraints, 200 kcal·mol$^{-1}$·rad$^{-2}$ for dihedral angle restraints, 50 kcal·mol$^{-1}$·Å$^{-2}$ for hydrogen bond-based distance restraints, 30 kcal·mol$^{-1}$·Å$^{-2}$ for planarity restraints, and 10 kcal·mol$^{-1}$·rad$^{-2}$ for chirality restraints. The system was equilibrated at 300 K for 5 ps and subsequently heated to 1000 K for 10 ps. This temperature was maintained for the next 30 ps. The system was then cooled to 100 K and finally to 0 K within the next 45 ps and 10 ps, respectively. Simulated annealing was carried out on 100 starting structures to yield twenty converged structures. For further refinement in water, ten lowest-energy conformations were selected. The system was neutralized by adding potassium ions. In addition, two potassium ions were placed within the inner channel of the G4 core between the eight O6 atoms of two adjacent G-tetrad layers. For hydrating the system, TIP3P water was added in a 10 Å truncated octahedral box. The simulation started with 500 steps of steepest descent followed by 500 steps of conjugate gradient minimization. For initial equilibration, the DNA was fixed with a force constant of 25 kcal·mol$^{-1}$·Å$^{-2}$. The system was then heated from 100 to 300 K in 10 ps under constant volume, and equilibration continued at 1 atm with energy restraints reduced from 5 to 0.5 kcal·mol$^{-1}$·Å$^{-2}$. Finally, the simulation was run for 4 ns at 1 atm and 300 K using NMR-derived distance, Hoogsteen hydrogen bond, and planarity restraints. The restraint energies were set to 15 kcal·mol$^{-1}$·Å$^{-2}$ for NOE-based distance restraints, to 25 kcal·mol$^{-1}$·Å$^{-2}$ for hydrogen bond-based distance restraints, and to 5 kcal·mol$^{-1}$·Å$^{-2}$ for planarity restraints. The trajectory was averaged over the last 500 ps and minimized in vacuum to obtain the final ten lowest-energy structures.

### 2. Biophysical Analysis of Ligand interaction with *ORAI1-Pu*

#### 2.1. Isothermal titration calorimetry

The thermodynamic interaction profile between ligand and putative DNA was studied using an iTC200 microcalorimeter at 25˚C. The *ORAI1-Pu* synthetic oligonucleotide and the ligands TMPyP4 and BRACO-19 were diluted in potassium phosphate buffer (10 mM) to the required concentration (1:50), such that the titration reached a saturation point. They were degassed in a vacuum for 10 min to ensure the removal of bubbles. The DNA was then added to the cell, and the ligand (1mM) to the syringe was set to eject 2µl per every 120 sec into the cell containing *ORAI1-Pu* (20µM). A control experiment was also conducted in parallel with the ligand (1mM) titrated into the buffer without oligonucleotide to subtract the heat of dilution from the ligand–quadruplex binding experiment before curve fitting. The number of injections was set to 20 to reach binding saturation. By integrating the area of peaks with Nano-analyze build-in Origin 7.0 software, the heat of reaction per injection was determined, and the enthalpy of binding ΔH°, the stoichiometry of reaction (n), and the dissociation constant $K_d$ were extracted from the best fit using an independent binding site model with $\chi^2$ values inspected for the best curve fit averaged over two independent experiments. (Figure S8 and Table S3)

#### 2.2 CD melting experiments

For CD melting experiments, *ORAI1-Pu* (5 µM) and the complexes (*ORAI1-Pu*: BRACO-19/TMPyP4 with a 1:2 DNA: ligand molar ratio) were dissolved in 10 mM potassium phosphate buffer, pH 7.0. Ellipticities at 263 nm were recorded from 25 °C to 90 °C in 5 °C increments. The heating rate was 0.2 °C min$^{-1}$ and the bandwidth was 1 nm. Melting temperatures were determined from the first derivative of the melting curve and averaged over two independent experiments. (Figure S9)

#### 2.3. NMR titration experiments

*ORAI1-Pu* oligonucleotides were annealed in KP buffer (10 mM) by heating at 95°C for 5 min and subsequent gradual cooling. The sample was then dissolved in a 90% $H_20$/10% $D_2O$ solvent to a concentration of ~350 µM and its 1D proton NMR spectrum was acquired with the *"zgesgp"* pulse sequence and processed in TopSpin3.2. (Figure S9)

### 3. Supplementary Figures (S1-S11)

A)

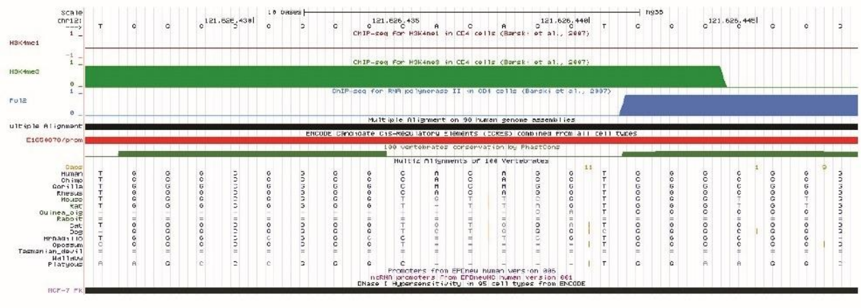

B)

QGRS sequences found (overlaps not included)

| Position | Length | QGRS | G-Score |
|---|---|---|---|
| 163 | 30 | GGGATATGGGCCTCAGGAGGCGCCCAGAGG | 19 |
| 215 | 30 | GGAGTCCAAGCTCCAGGAGCAGGGCCACGG | 12 |
| 255 | 12 | GGAGAGGGGCGG | 18 |
| 272 | 21 | GGAGCCGGAGCGGGCAGCCGG | 20 |
| 310 | 12 | GGCGGGCGGCGG | 20 |
| 341 | 21 | GGTGCCGTCGGAGCGGGCGGG | 17 |
| 397 | 23 | GGGCGGGGCACAGGTGGGCGGGG | 37 |
| 438 | 19 | GGGCGGCGGCCAATCGCGG | 14 |
| 466 | 26 | GGGGTCCAGGCCCCGGGGATCCGAGG | 21 |

**Figure S1**: A) UCSF browser data for the putative G4 region. The region is conserved in primates and shows variability in the loop region in other mammals. Session link: https://genome-euro.ucsc.edu/s/oishika/hg38_Orai1%2DPu_region. B) QGRS predictor putative G4 motifs within -499bp from TSS: *ORAI1-Pu* has the highest G4 score and has been further selected for our study.

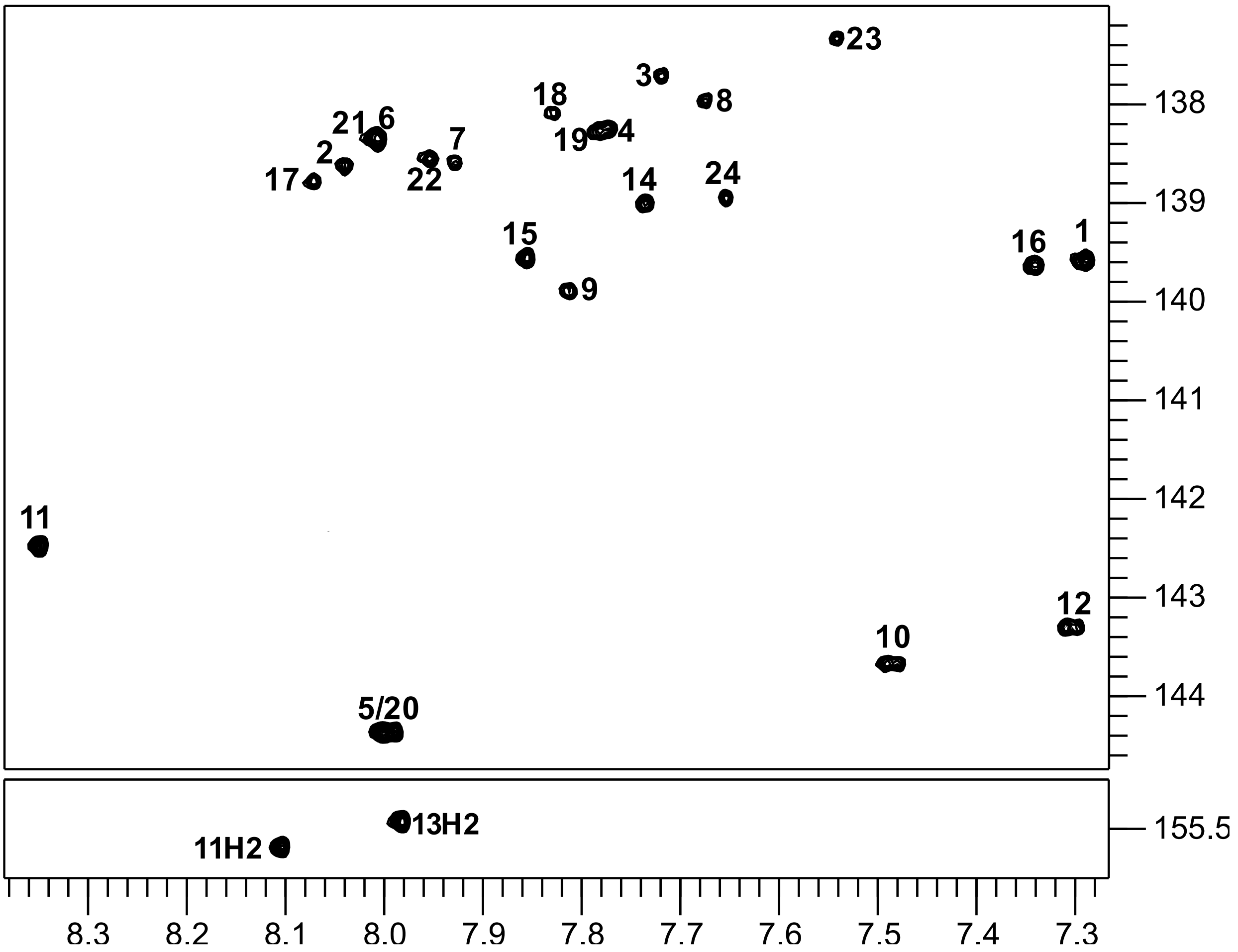


**Figure S2:** $^{1}H$-$^{13}C$ HSQC spectrum of *ORAI1-Pu* showing H8/H6($\omega_2$)-C8/C6($\omega_1$) (top) and adenine H2($\omega_2$)-C2($\omega_1$) correlations (bottom). NMR spectra were acquired in a 10 mM potassium phosphate buffer, pH 7.0, at 40 °C.

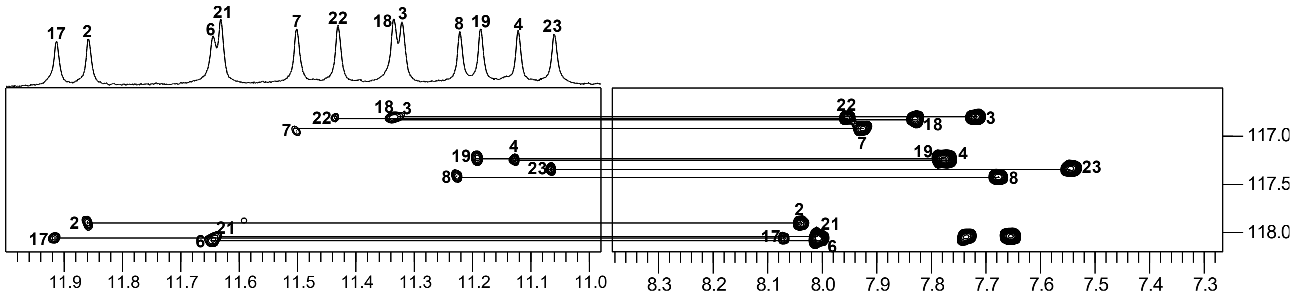


**Figure S3:** $^{1}$H-$^{13}$C HMBC spectrum of *ORAI1-Pu* showing correlations between guanine H1($\omega_2$) and H8($\omega_2$) resonances through their long-range coupling to $^{13}$C5($\omega_1$) at natural abundance. NMR spectra were acquired in a 10 mM potassium phosphate buffer, pH 7.0, at 40 °C.

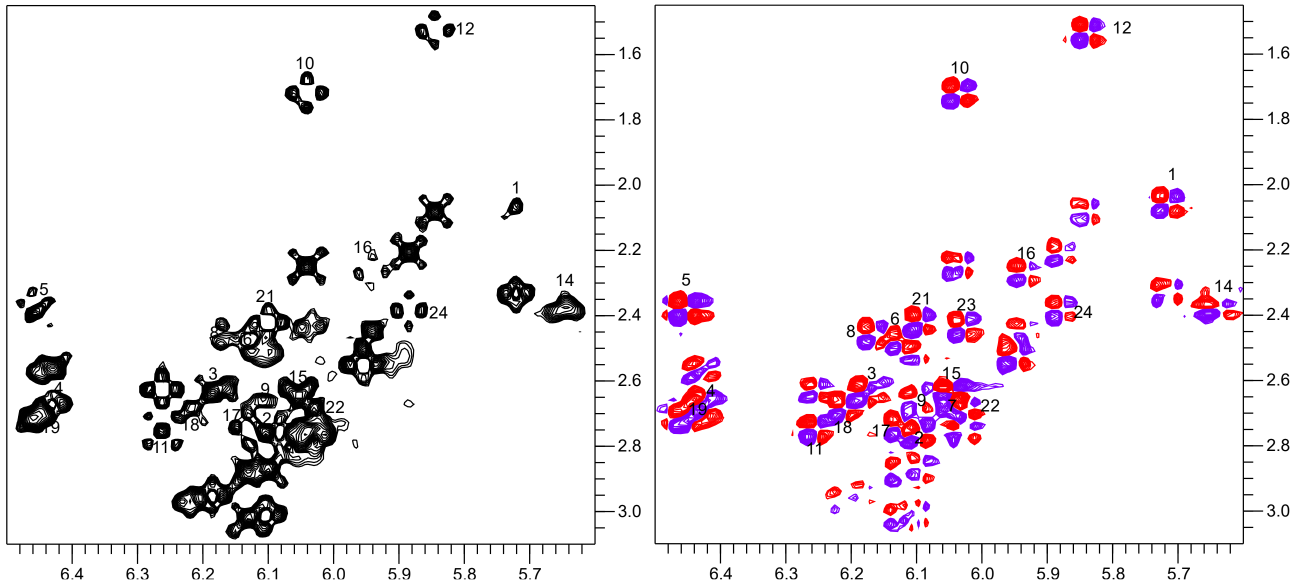


**Figure S4:** Stereospecific assignments of H2'/H2" resonances. H1'($\omega_2$)-H2'/H2"($\omega_1$) NOESY spectral region acquired at short mixing time (80 ms) (left) and H1'($\omega_2$)-H2'/H2"($\omega_1$) DQF-COSY spectral region showing H1'-H2' and H1'-H2" intra-residual crosspeaks for the *ORAI1-Pu* sequence at 40°C.

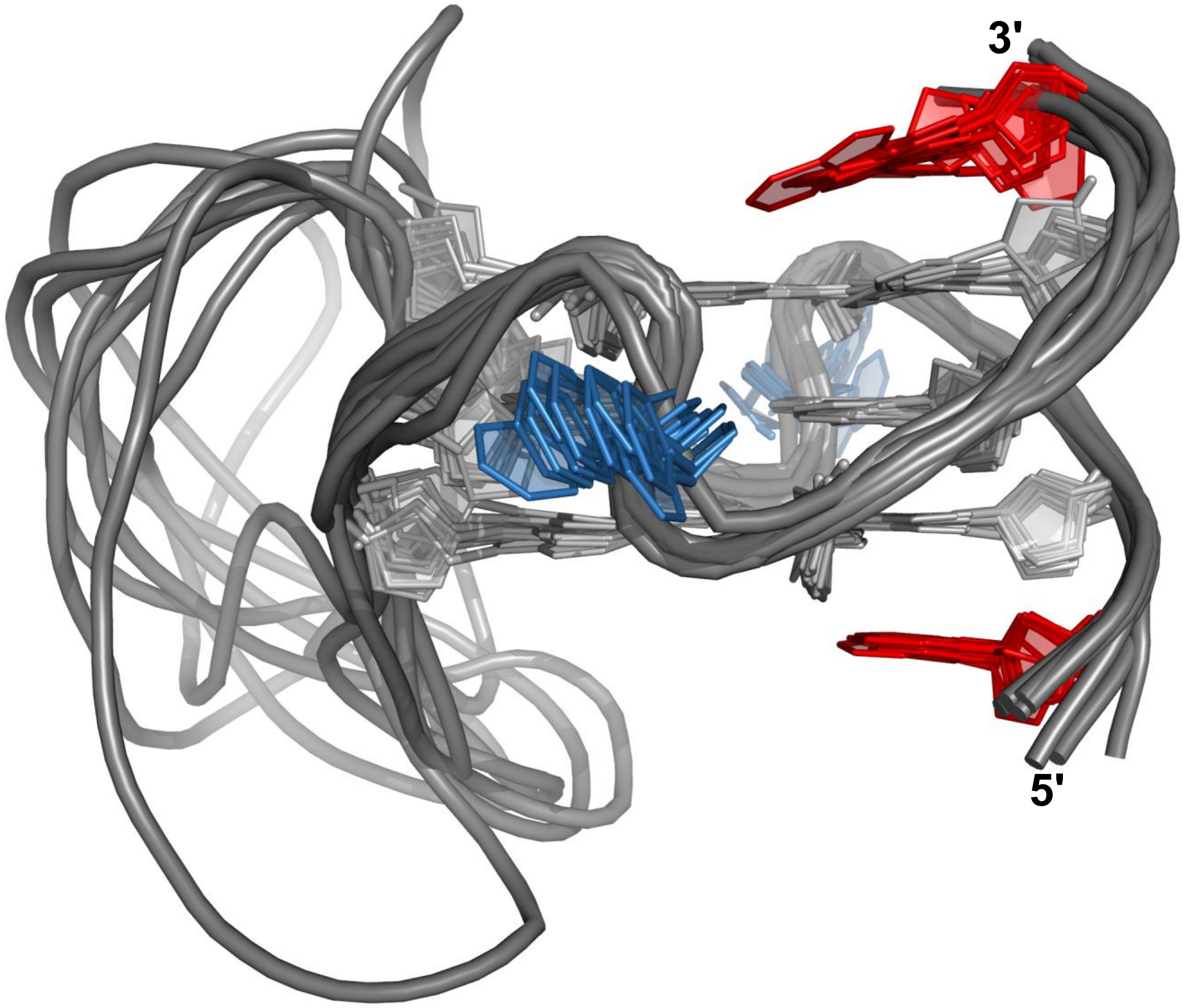


**Figure S5:** Superposition of ten lowest-energy structures of *ORAI1-Pu*; residues of the second propeller loop have been omitted for clarity. *Anti*-G-residues are colored grey, loop residues are colored blue, and overhang residues are colored red.

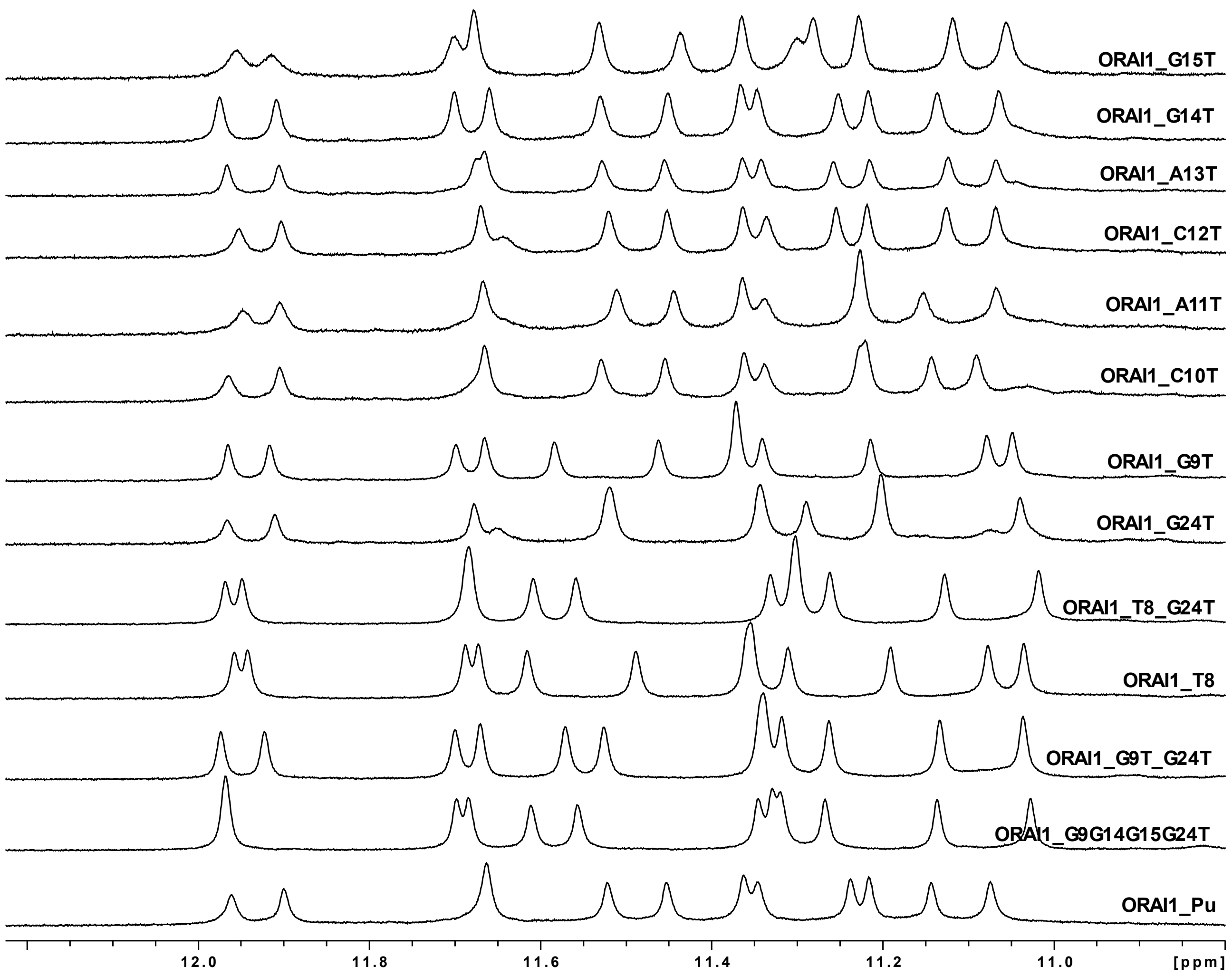


**Figure S6:** Imino proton spectral region of *ORAI1-Pu* mutants. NMR spectra were acquired in a 10 mM potassium phosphate buffer, pH 7.0, at 25°C.

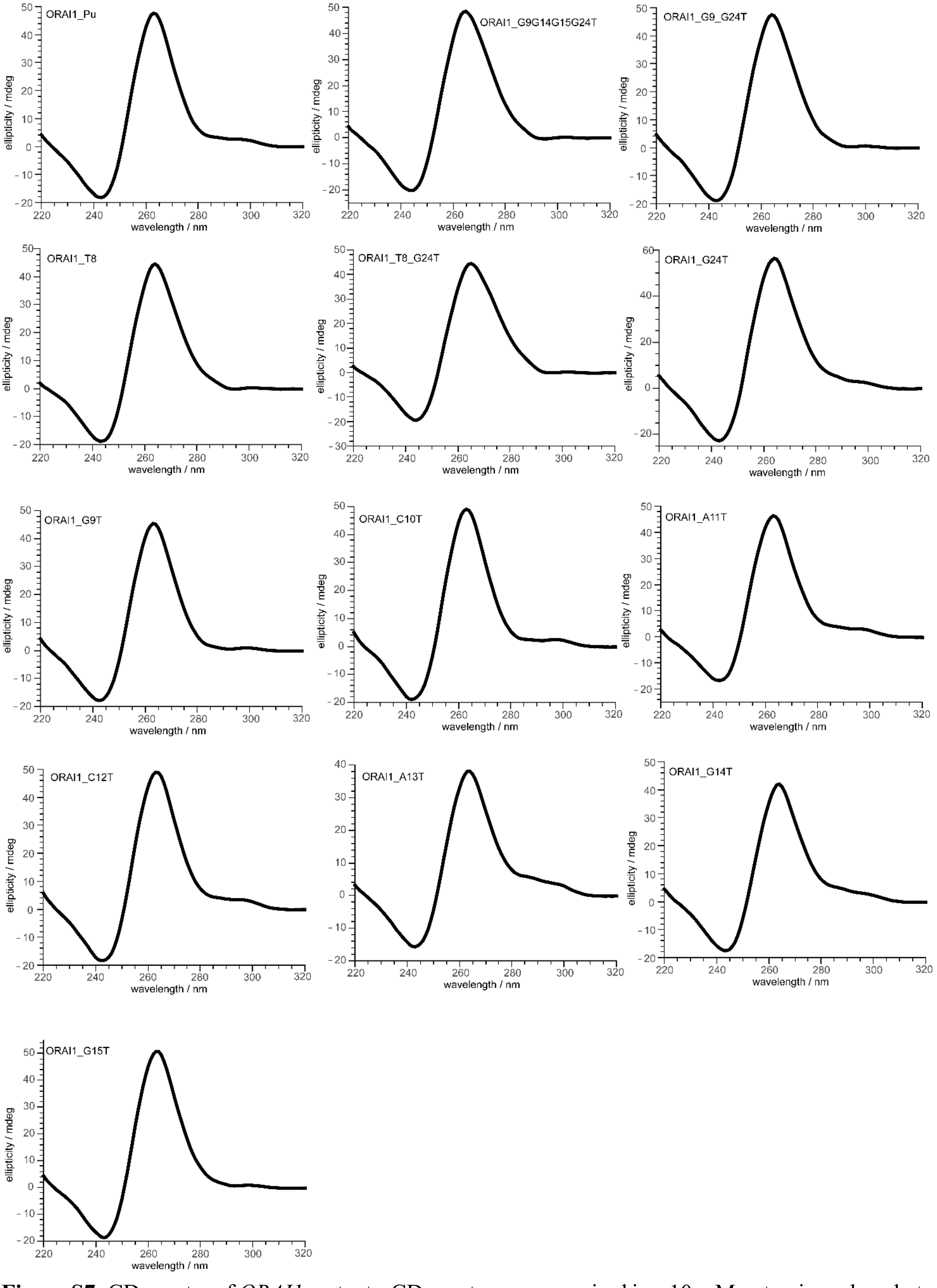


**Figure S7:** CD spectra of *ORAI1* mutants. CD spectra were acquired in a 10 mM potassium phosphate buffer, pH 7.0, at 20°C.

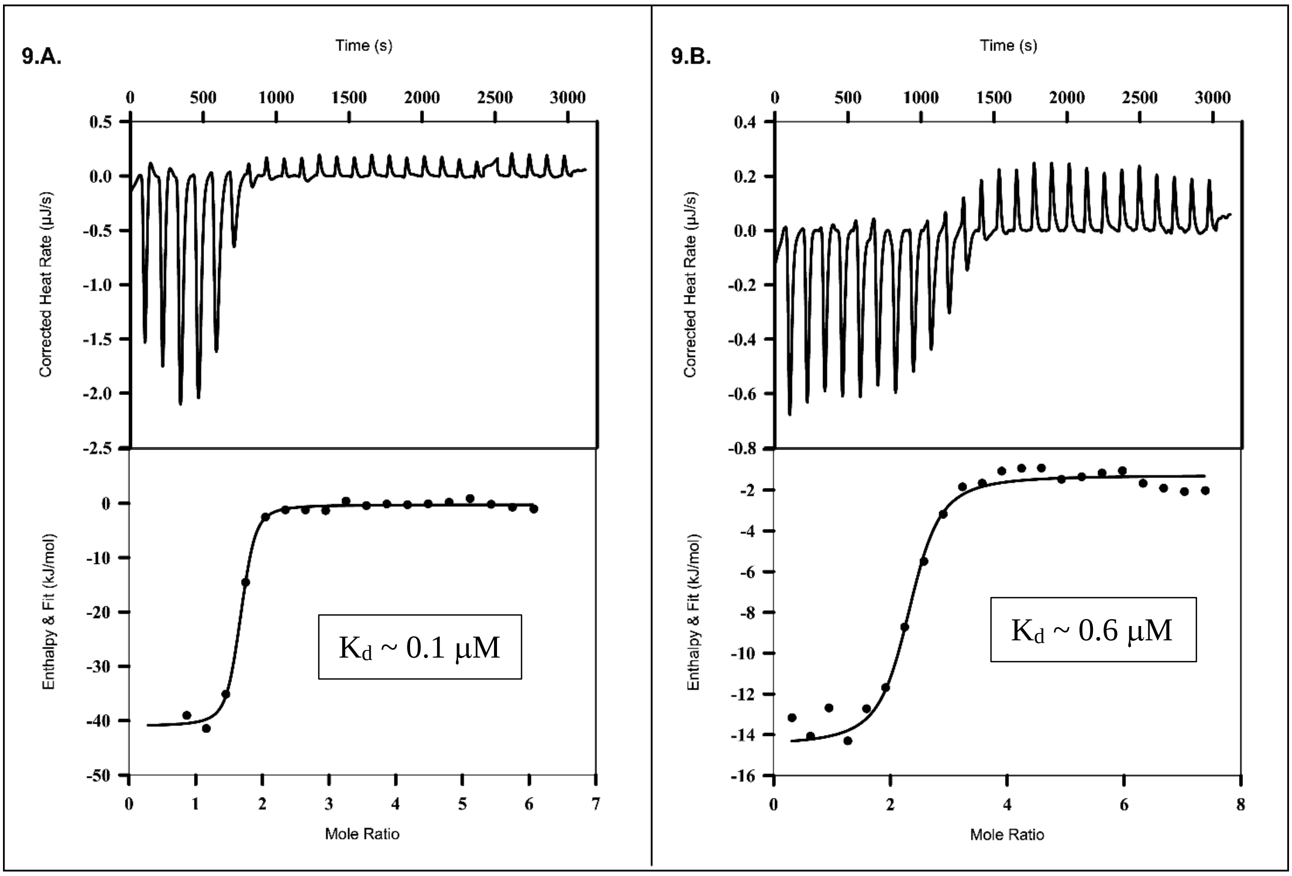


**Figure S8**: ITC profiles for the interaction of TMPyP4 (9.A.) and BRACO-19 (9.B.) with *ORAI1-Pu*. Detailed thermodynamic parameters are given in the Supplementary Table S3.

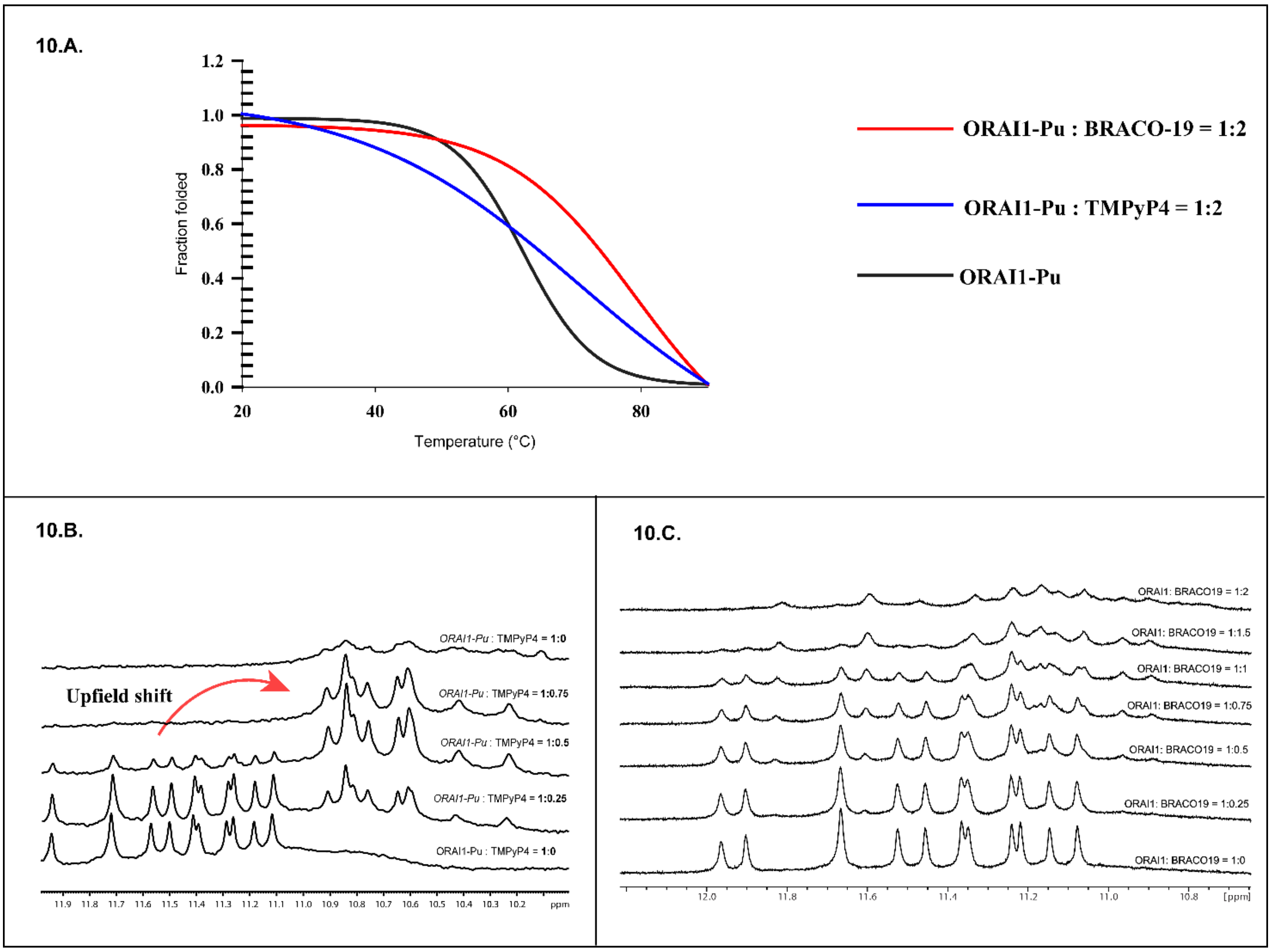


**Figure S9**: **(A)** CD melting curves in KP buffer (10 mM); *ORAI1-Pu* ($T_m$ = 61.4 °C), *ORAI1-Pu*: TmPyP4 ($T_m$ = c), *ORAI1-Pu*: BRACO-19 ($T_m$= 80 ± 2 °C). (**B**) and (**C**) 1D imino proton NMR spectral region with increasing TMPyP4 **(B)** and BRACO-19 **(C)** concentration.

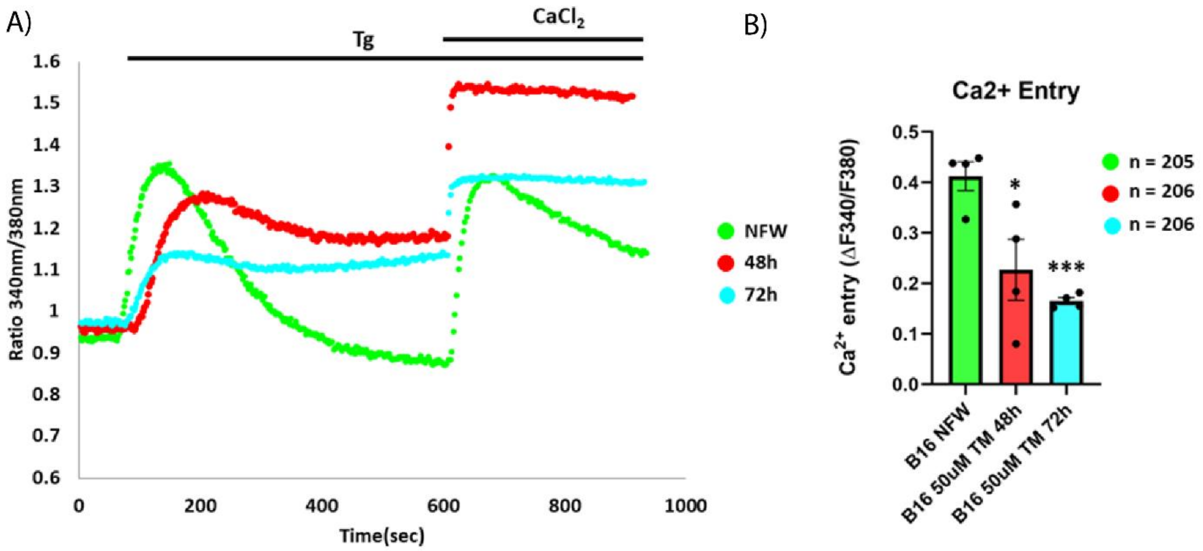


**Figure S10: TMPyP4 decreases *ORAI1* mediated SOCE in B16 cells.** (A) Representative $Ca^{2+}$ imaging traces for control nuclease-free water i.e. NFW and TMPyP4 pre-treatment for 48-72hrs. Cells were stimulated with 2µM thapsigargin (Tg) in $Ca^{2+}$ free buffer followed by restoration with 1mM extracellular $Ca^{2+}$. (B) The extent of SOCE was calculated from 205 NFW and 206 TMPyP4 treated B16 cells, which were imaged from 3 independent experiments ("n = x" where "x" denotes total number of cells imaged). Data presented are mean ± S.E.M. For statistical analysis, unpaired student's t test was performed. Here, * means $p < 0.05$, *** means $p < 0.001$.

### 4. Supplementary Tables (S1-S5)

**Table S1:** $^{1}$H and $^{13}$C chemical shifts (ppm) of *ORAI1-Pu* at 40 ºC.

| Residue | H8/H6 | H1/H3 | H1' | H2'/H2'' | H3' | H5/H2/Me | C8/C6 | C2 |
|---|---|---|---|---|---|---|---|---|
| T1 | 7.29 | - | 5.72 | 2.06/2.33 | 4.65 | 1.35 | 139.58 | - |
| G2 | 8.04 | 11.86 | 6.10 | 2.77/3.01 | 4.99 | - | 138.62 | - |
| G3 | 7.72 | 11.32 | 6.18 | 2.63/2.95 | 5.00 | - | 137.72 | - |
| G4 | 7.77 | 11.12 | 6.43 | 2.67/ 2.56 | 5.08 | - | 138.25 | - |
| C5 | 8.00 | - | 6.46 | 2.38/2.71 | 5.07 | 6.16 | 144.38 | - |
| G6 | 8.01 | 11.65 | 6.13 | 2.48/2.88 | 5.12 | - | 138.33 | - |
| G7 | 7.93 | 11.50 | 6.05 | 2.67/ 2.76 | 5.03 | - | 138.59 | - |
| G8 | 7.68 | 11.22 | 6.17 | 2.46/2.63 | 4.90 | - | 137.96 | - |
| G9 | 7.82 | - | 6.11 | 2.66/2.51 | 4.89 | - | 139.88 | - |
| C10 | 7.49 | - | 6.04 | 1.73/2.25 | 4.73 | 5.88 | 143.67 | - |
| A11 | 8.35 | - | 6.26 | 2.75/2.63 | 4.92 | 8.11 | 142.48 | 155.69 |
| C12 | 7.31 | - | 5.84 | 1.54/2.08 | - | 5.72 | 143.30 | - |
| A13 | 8.06 | - | 5.96 | 2.51/2.54 | 4.83 | 7.98 | 142.22 | 155.43 |
| G14 | 7.74 | - | 5.64 | 2.38/2.39 | 4.86 | - | 139.01 | - |
| G15 | 7.86 | - | 6.05 | 2.62/2.64 | 4.90 | - | 139.56 | - |
| T16 | 7.34 | - | 5.94 | 2.27/2.45 | 4.83 | 1.52 | 139.63 | - |
| G17 | 8.07 | 11.91 | 6.13 | 2.74/ 3.02 | 4.95 | - | 138.78 | - |
| G18 | 7.83 | 11.34 | 6.22 | 2.68/2.97 | 5.07 | - | 138.10 | - |
| G19 | 7.79 | 11.19 | 6.44 | 2.69/2.56 | 5.09 | - | 138.28 | - |
| C20 | | | | | | | | |
| G21 | 8.01 | 11.63 | 6.10 | 2.42/2.87 | 5.14 | - | 138.33 | - |
| G22 | 7.96 | 11.43 | 6.03 | 2.68/2.76 | 5.05 | - | 138.54 | - |
| G23 | 7.54 | 11.06 | 6.03 | 2.43/2.45 | 4.94 | - | 137.34 | - |
| G24 | 7.66 | | 5.88 | 2.39/2.21 | | | 138.94 | |

**Table S2: *ORAI1-Pu* and mutant sequences with their UV melting temperatures $T_{m}$.**

| Name | Sequence | $T_{m}$ (°C) |
|---|---|---|
| ORAI1_Pu | TGGGCGGGGCACAGGTGGGCGGGG | 61.4±0.01 |
| ORAI1_G9G14G15G24T | TGGGCGGGTCACATTTGGGCGGGT | 54.8±0.1 |

| | | |
|---|---|---|
| ORAI1_G9T_G24T | TGGGCGGGTCACAGGTGGGCGGGT | 57.6±0.5 |
| ORAI1_T8 | TGGGCGGGTTTTTTTTGGGCGGGG | 56.4±0.4 |
| ORAI1_T8_G24T | TGGGCGGGTTTTTTTTGGGCGGGT | 53.0±0.4 |
| ORAI1_G24T | TGGGCGGGGCACAGGTGGGCGGGT | 58.4±0 |
| ORAI1_G9T | TGGGCGGGTCACAGGTGGGCGGGG | 61.1±0.2 |
| ORAI1_C10T | TGGGCGGGGTACAGGTGGGCGGGG | 60.9±0.05 |
| ORAI1_A11T | TGGGCGGGGCTCAGGTGGGCGGGG | 61.2±0.4 |
| ORAI1_C12T | TGGGCGGGGCATAGGTGGGCGGGG | 61.1±0.2 |
| ORAI1_A13T | TGGGCGGGGCACTGGTGGGCGGGG | 61.1±0.5 |
| ORAI1_G14T | TGGGCGGGGCACATGTGGGCGGGG | 60.8±0.02 |
| ORAI1_G15T | TGGGCGGGGCACAGTTGGGCGGGG | 61.1±0.2 |

**Table S3: Thermodynamic Parameters by ITC.**

| Ligand | Variables | |
|---|---|---|
| TMPyP4 (independent binding model) | n | 1.534 ± 0.22 |
| | ΔH° (kJ/mol) | -40.77 ± 1.5888 |
| | ΔS° (J/mol -K) | -3.314 |
| | ΔG° (kJ/mol) | -39.78 |
| | $K_d$ (nM) | 107.4 ± 55.14 |
| BRACO-19 (independent binding model) | n | 2.192 ±0.081 |
| | ΔH° (kJ/mol) | -13.25 ±1.313 |
| | ΔS° (J/mol -K) | 74.14 |
| | ΔG° (kJ/mol) | -35.344 |
| | $K_d$ (nM) | 638.6 ± 340.6 |

**Table S4: Primers used for q-PCR analysis and ChIP assay**

| Primers | Sequence | Melting Temp |
|---|---|---|
| ***ORAI1*-Forward** | CAGAGTTACTCCGAGGTGATGAG | 59°C |
| ***ORAI1*-Reverse** | GAGAGCAGAGAGGAGGTCC | 60°C |
| **18S rRNA-Forward** | CGGACAGGATTGACAGATTGATAGC | 59°C |
| **18S rRNA-Reverse** | TGCCAGAGTCTCGTTCGTTATCG | 61°C |
| ***ORAI1*-Pu-Forward (ChIP)** | CAGAGTTACTCCGAGGTGATGAG | 59°C |
| ***ORAI1*-Pu-Reverse (ChIP)** | GAGAGCAGAGAGGAGGTCC | 60°C |

**Table S5: Antibodies used for ChIP assay**

| Name of antibody used | Dilution used | Company | Catalogue No. |
|---|---|---|---|
| **RNA Pol** | 1:100 | Thermo-Fischer (Included in Pierce ChIP kit) | 1862739 |
| **IgG** | 1:500 | Thermo-Fischer (Included in Pierce ChIP kit) | 1862739 |
| **E2F1** | 1:500 | Invitrogen | KH95 |
| **c-Myc** | 1:500 | Santa Cruz Biotechnology | 9E10 |
| **SNAI2/Slug (A-7)** | 1:500 | Santa Cruz Biotechnology | SC-166476 |
| **BG4** | 1:250 | Sigma-Aldrich | MABE917 |
| **ZEB1** | 1:500 | Cell Signaling Technology | CST-70512T |
